# A Recurring Chemogenetic Switch for Chimeric Antigen Receptor T Cells

**DOI:** 10.1101/2021.08.23.457355

**Authors:** Wenyue Cao, Zhi Z. Geng, Na Wang, Quan Pan, Shaodong Guo, Shiqing Xu, Jianfeng Zhou, Wenshe Ray Liu

## Abstract

As a revolutionary cancer treatment, the chimeric antigen receptor (CAR) T cell therapy suffers from complications such as cytokine release syndromes and T cell exhaustion. Their mitigation desires controllable activation of CAR-T cells that is achievable through regulatory display of CARs on the T cell surface. By embedding the hepatitis C virus NS3 protease (HCV-NS3) in an anti-CD19 CAR between the anti-CD19 single-chain variable fragment (scFv) and the hinge domain, we showed that the display of anti-CD19 scFv on CAR-T cells was positively correlated to the presence of a clinical HCV-NS3 inhibitor asunaprevir (ASV). This novel CAR design that allows the display of anti- CD19 scFv on the T cell surface in the presence of ASV and its removal in the absence of ASV effectuates a practically recurring chemical switch for CAR-T cells. We demonstrated that the intact CAR on T cells was repeatedly turn on and off by controlling the presence of ASV. The dose dependent manner of the intact CAR display on T cells with regard to the ASV concentration enables delicate modulation of CAR-T cell activation during cancer treatment. In a mouse model, we showed different treatment prospects when ASV was provided at different doses to mice that were infused with both human CD19^+^ lymphoma and the switchable CAR-T cells.

## Introduction

Chimeric antigen receptors (CARs) refer to engineered T cell receptors that recognize specific antigens on the cancer cell surface and subsequently direct CAR-T cells selectively to cancer cells for their elimination. The advent of the CAR-T cell therapy technique has been revolutionizing clinical intervention of cancer especially for hematological malignancies^1–5^. In clinical practice, T cells are generally isolated from the patient’s own blood, stimulated for proliferation, and then transduced with a gene encoding an engineered CAR via virus or non-virus methods. The CAR structure contains typically an extracellular antigen-binding domain and intracellular signaling domains for an enhanced response to antigen recognition. After engineered CAR-T cells are infused back into the patient, they proliferate robustly to mount a potent immune response against malignant cells for their elimination^6–8^. Crowned as live drugs, CAR-T cells have demonstrated success in the elimination of hematopoietic tumors including different types of leukemia, lymphoma, and myeloma. CAR-T cell therapeutics that have been approved by the U.S. Food and Drug Administration for clinical uses include Kymriah, Yescarta and Tecartus and a lot more are on clinical trials^9–12^. Although extremely powerful, the CAR-T cell therapy has potentially serious safety concerns. Uncontrollable activation of CAR-T cells can trigger life-threatening adverse events including clinically significant release of proinflammatory cytokines termed as cytokine release syndromes, encephalopathy, multi-organ failure, and eventual death^13–16^. Risks for severe cellular toxicity present a key challenge to the effective administration of CAR-T cells on a routine basis. Another effect of uncontrollable activation of CAR-T cells is T cell exhaustion in which chronical antigen stimulation triggers a progressive loss of T cell functions^17^.

To mitigate limitations related to uncontrollable CAR-T cell activation, temporal and spatial control of CAR-T activity is desired. To achieve it, a number of approaches have been developed and tested. Proapoptotic safety switches, like suicides genes, have been designed. However, suicide switches do not provide control over CAR-T cell activation or expansion whereas they result in irreversible elimination of therapeutic CAR-T cells^18, 19^. Another strategy to achieve controllable CAR-T cell activation is to use small molecules or protein adapters to gate cellular functions^20–24^. Bifunctional antibodies that serve as a bridge between CAR-T and tumor cells have also been designed as switches^25^. Small molecules may also be used as inducers for CAR expression. Both small molecules and protein adapters can be administered at varying doses, which have an advantage on precise regulation over the activity of therapeutic CAR-T cells. However, many of these small molecules are not clinically approved medications^20, 26, 27^. Their combined uses with CAR-T cells pose significant challenges in clearing safety concerns in patients. Most current switches implement controllable display of CARs on the CAR-T cell surface when the switch is turned on. When the switch is off, T cells with no functional CARs are produced; however, the off condition has typically no effect on existing functional CARs on the T cell surface. A switch that allows recurring turning-on and off of CARs on same CAR-T cells has not been devised. In this paper, we would like to report the development of such a switch controlled by a small molecule Asunaprevir (ASV) (Fig. 1).

**Fig. 1:**
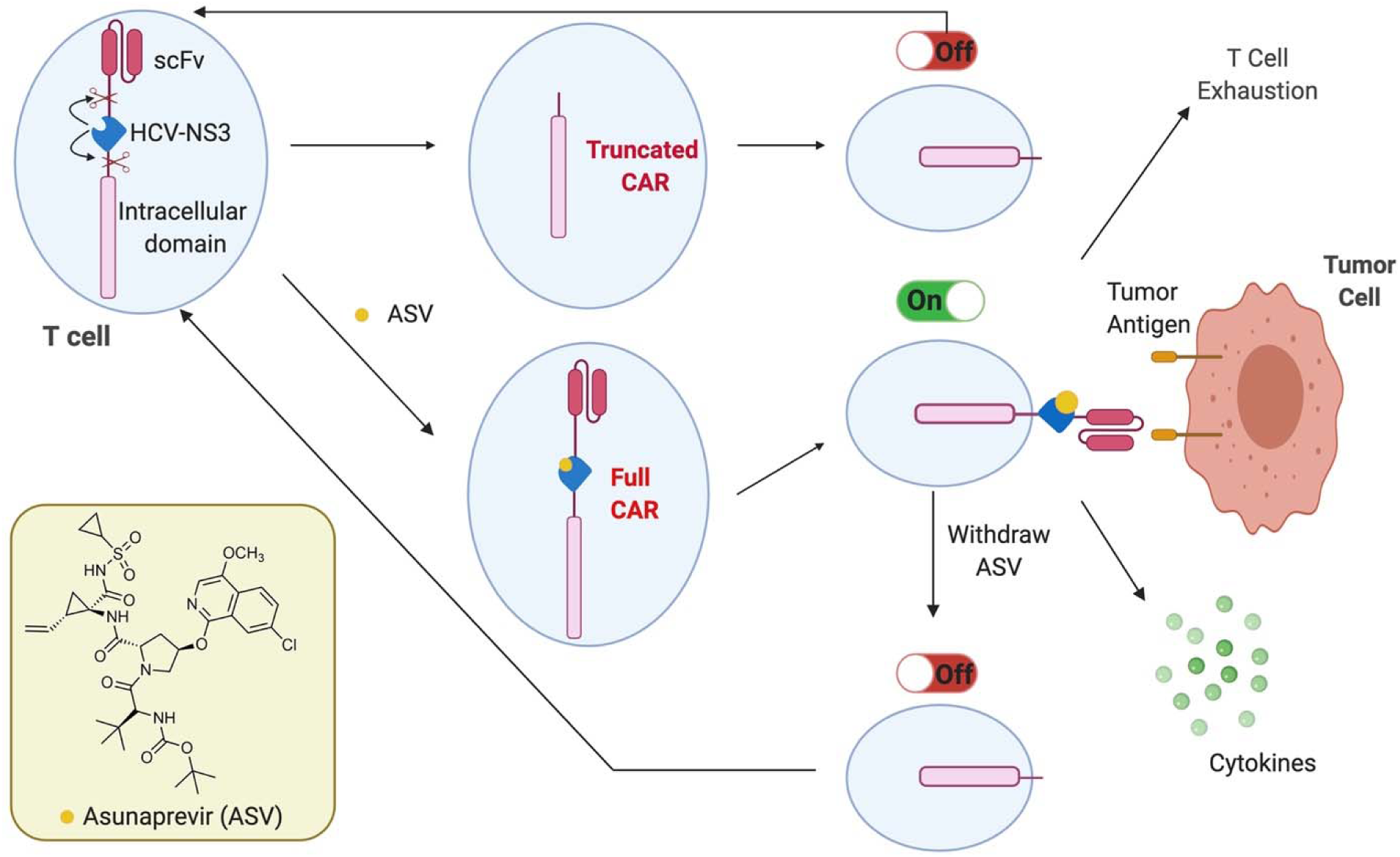
**Graphic illustration of a recurring chemogenetic switch that uses asunaprevir (ASV) in coordination with the hepatitis C virus NS3 protease (HCV-NS3) to regulate CAR presentation on the T cell surface. In the absence of ASV, T cells undergo proliferation without turning active while the presence of ASV triggers full CAR display to activate T cells for immunogenic elimination of tumor cells along with cytokine release and potential T cell exhaustion. Removal of ASV switches off active CAR-T cells by cleaving the displayed CAR.**

## Results and Discussion

### Design, preparation and characterization of a recurring chemogenetic CAR switch

To achieve both on and off effects of a switch in a same CAR-T cell, we set our sights on chemogenetic control of protein functions for our switch design. From the perspectives of clinical use of a switchable CAR-T cell therapeutic, the molecule used in the chemogenetic control needs to pass all safety requirements. As a hybrid therapeutic with both a small molecule and CAR-T cells, a practical exercise is to use a control molecule that has already been approved for clinical use in human patients. The use of an approved drug will avoid potential safety complications during clinical trials and expedite the clinical approval of a switchable CAR-T therapeutic since one component in the hybrid therapeutic has been cleared with all safety concerns. We deemed that the hepatitis C virus nonstructural protein 3 protease (HCV-NS3) and its clinically approved inhibitors such as ASV fulfill the requirements necessary as a safe chemogenetic switch^28, 29^. HCV-NS3 is a polypeptide domain of a much larger HCV polypeptide translate that undergoes autoproteolysis. The autoproteolysis nature of HCV-NS3 that hydrolyses at its both ends and its inhibition by ASV have been previously used to develop stabilizable polypeptide linkages to control proteasome-based degradation of a protein target^30–32^. In order to build a chemogenetic switch based on HCV-NS3 that responds to ASV positively and degrade a displayed CAR on CAR-T cells in the absence of ASV to make the switch a practically recurring one, we followed a CAR structure design shown in Fig. 2a. We first constructed a vector for expressing a standard anti-human CD19 CAR (CAR19). The structure was based on the third-generation CAR design that contained an anti-CD19 single-chain variable fragment (scFv) derived from a murine monoclonal antibody against human CD19 and with a same amino acid sequence as the one in the approved CAR-T therapeutic Kymriah^12^, a CD8 hinge domain, a transmembrane (TM) domain, two costimulatory domains CD28 and 4-1BB for a strong antigen response, and an intracellular CD3ζ tail for homodimer formation. DNA encoding this design was cloned into a pLVX-EF1α vector to afford plasmid pLVX-EF1α-CAR19. Along with two other plasmids psPAX2 and PMD2.G, we used the three vectors to transfect HEK 293T/17 cells for the production of lentivirus particles that we used further to transduce human T cells isolated from leukocytes of human donors to produce standard CAR19 T cells. To install a recurring chemogenetic switch to CAR19 for effective removal of anti-CD19 scFv in the absence of ASV, we cloned HCV-NS3 between anti-CD19 scFv and the hinge domain in plasmid pLVX-EF1α CAR19 to afford plasmid pLVX-EF1α sCAR19 for the expression of a switchable CAR19 (sCAR19 in Fig. 1a). In sCAR19, a T54A mutation was introduced to HCV-NS3 to improve its sensitivity toward ASV and two HCV-NS3 cleavage sites were inserted, one between anti-CD19 scFv and HCV-NS3 and the other between the hinge and the TM domain. To improve the autoproteolysis activity of HCV-NS3, we followed a previous design by fusing an HCV NS4 cofactor fragment at the HCV-NS3 *N*-terminal side^31^. Plasmid pLVX-EF1 α -CAR19 was then used to generate sCAR19 T cells by following the same procedure described for the construction of CAR19 T cells.

**Fig. 2:**
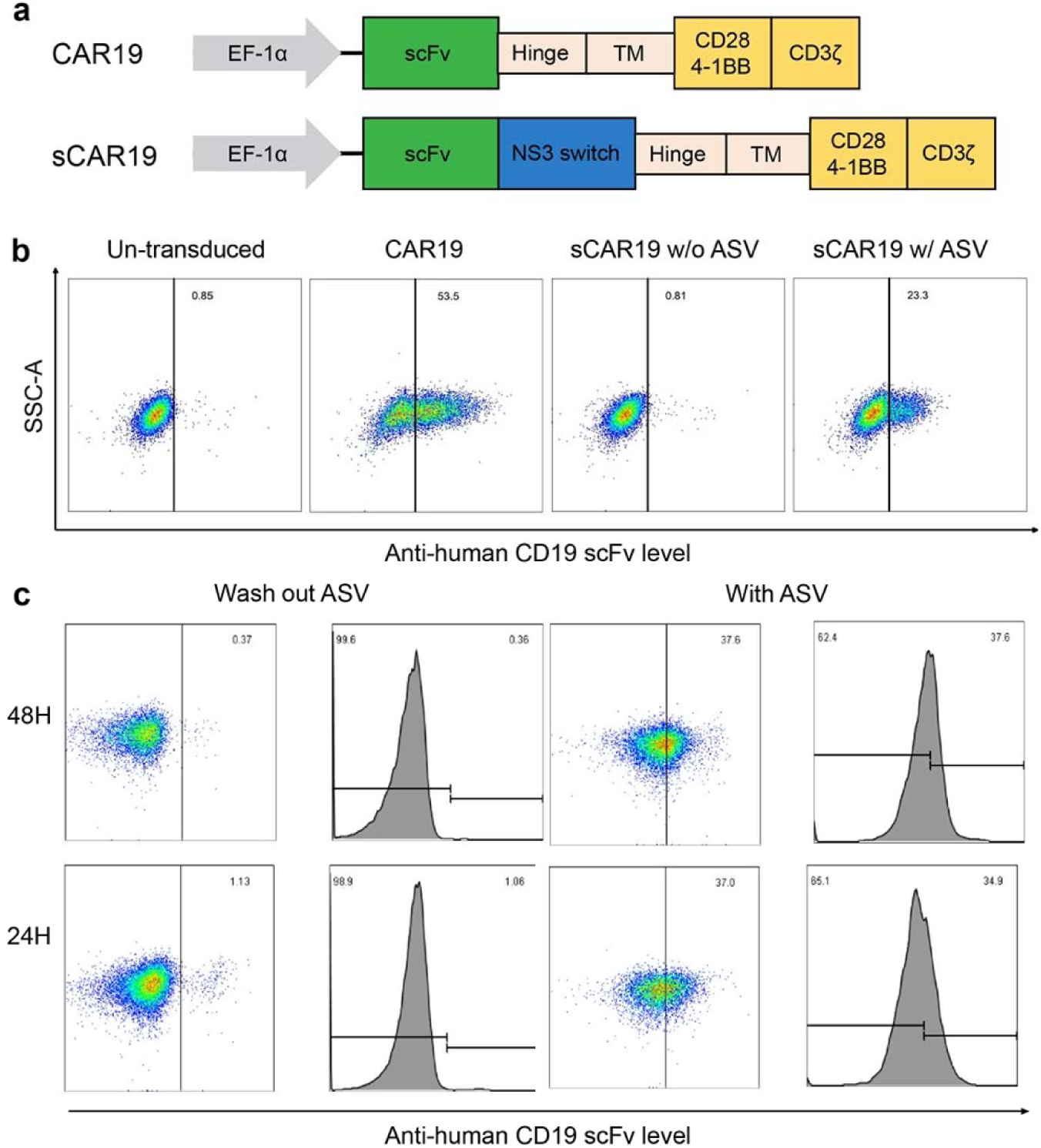
**The recurring chemogenetic switch demonstrates both on and off effects in the regulation of CAR display on the T cell surface. (a) Schematic representation of expression vector designs for a standard CAR19 and a switchable CAR19 (sCAR19) containing an HCV-NS3-based recurring switch. Both CARs are under control of an EF1**α promoter. (b) Density plots showing distinct anti-human CD19 scFv presentation on un-transduced T cells, CAR19 T cells, and sCAR19 T cells cultured under two conditions, one with a DMSO vehicle and the other with 1 μM ASV. The display of anti-human CD19 scFv was determined by flow cytometry using Alexa Fluor 647-anti-mouse F(ab)^2^ antibody. sCAR19 T cells were cultured with the vehicle or ASV for 24 h before the flow cytometry analysis. (c) The cleavage of anti-human CD19 scFv from sCAR19 T cells that were cultured originally with 1 μM ASV to display full-length sCAR19 and then with ASV withdrawn for 24 and 48 h. sCAR19 T cells that were cultured continuously with 1 μM ASV were used as controls. Alexa Fluor 647-anti-mouse F(ab)^2^ antibody was used to label cells for the flow cytometry analysis.

Unlike un-transduced T cells that expressed no CAR19 and the standard CAR19 T cells that displayed constitutively CAR19 on the cell surface, sCAR19 T cells displayed full-length sCAR19 with a positive correlation to the presence of ASV in the growth media (Fig. 2b). The absence of ASV led to close to no full-length sCAR19 that we could detect using Alexa Fluor 647-anti-mouse F(ab)^2^ antibody. However, the addition of 1 μM ASV to the growth media resulted in the display of full-length sCAR19 on the cell surface (24% of all sorted cells). When ASV was withdrawn from the media, the displayed full-length sCAR19 on sCAR19 T cells degraded gradually. After 24 h, cells maintained a very low but detectable CAR19 (1% of all sorted cells) and after 48 h, close to no cells had detectable CAR19 (Fig. 2c). A control experiment in which we cultured sCAR19 T cells in the continuous presence of 1 μM ASV showed no difference on displayed full-length sCAR19 in the same time range. Collectively, our data establish that the display of full-length sCAR19 on sCAR19 T cells can be recurrently regulated by the presence of ASV and withdrawing ASV can effectively lead to total cleavage of full-length sCAR19 on all sCAR19 T cells.

To evaluate whether the level of displayed full-length sCAR19 can be regulated by the dose of ASV, we cultured sCAR19 T cells in the presence of serial concentrations of ASV ranging from 10 nM to 5 μM for 10 h (Extended Data Fig. 1). The displayed full-length sCAR19 had its highest level at 5 μM ASV and its level gradually decreased when the ASV dose reduced (Extended Data Fig. 2). Normalization of the data indicated that displayed full-length sCAR19 at 10 nM ASV was equivalent to about 50% of that at 5 μM. However, almost no displayed full-length sCAR19 was detected when ASV was absent in the growth media. These data indicated that the designed recurring chemogenetic switch was highly sensitive to the presence of ASV. To better understand how the switch works along with time, we cultured sCAR19 T cells in the presence of different concentrations of ASV for different periods of time with 2 h intervals from 0 to 10 h as well as for 24 h. Cells sorted with anti-mouse F(ab)^2^ antibody indicated a steady increase of displayed full-length sCAR19 from 0 to 10 h and reached close to the plateau after 10 h (Extended Data Fig. 3). Interestingly the displayed full-length sCAR19 level at 5 μM ASV and after 10 h culturing was almost as same as the one at 100 nM ASV and after 24 h culturing, indicating a relatively low dose of ASV can achieve efficient display of full-length sCAR19 on the CAR-T cell surface.

### Effects of the recurring chemogenetic switch on CAR-T characteristics

HCV-NS3 is a protease. Its expression in CAR-T cells may lead to undesired phenotypical consequences. To diffuse this concern, we analyzed the effects of both the switch and ASV on the T cell subset distribution and apoptosis. We cultured switchable sCAR19 T cells with or without 1 μM ASV for 3 days and then sorted subsets of T cells using differently colored antibodies for CD4 and CD8. Cell proliferation and early and late apoptotic cells were also analyzed. We carried out same analysis with un-transduced T cells and unswitchable CAR19 T cells as controls. For sCAR19 T cells, providing 5 *μ*M ASV to their growth media did not alter noticeably the distribution between CD4^+^ and CD8^+^ T cells that maintained around 53% and 43% respectively of the total cells (Extended Data Fig. 4). This distribution was also not significantly different for that in un-transduced and CAR19 T cells. For sCAR19 T cells that were grown with and without 5 μM ASV, their early and late apoptotic cell levels were almost identically around 2-3% and 8-9% respectively of the total cells. Both un-transduced and CAR19 T cells also had an early apoptotic level around 2% and late apoptotic cells around 10% (Extended Data Fig. 5). Collectively, data presented here approve that the recurring chemogenetic switch and ASV do not significantly alter T cell characteristics such as CD4^+^/CD8^+^cell distribution and apoptotic cell rates in comparison to a standard CAR design.

We have also analyzed T cell activation in sCAR19 T cells that were grown in the presence of 5 μM ASV in comparison to un-transduced and CAR19 T cells. The constitutively expressed CD25 was detected in 82-85% of the total cells in all three cell types. Similarly, in both switchable sCAR19 and unswitchable CAR19 T cells, CD69, a T cell activation marker exhibited close to an identical detection level as in un-transduced T cells (Extended Data Fig. 6). No significant difference was found between the un-transduced and switchable sCART cells (P > 0.05).

### ASV-regulated cytotoxicity of switchable CAR-T cells *in vitro*

With demonstrated controllable T cell activation for sCAR19 T cells in the presence of ASV, we proceeded further to test ASV-regulated cytotoxic effects of sCAR19 T cells in killing CD19^+^ Raji cells that were derived from Burkitt’s lymphoma. sCAR19 T cells were cultured in the presence of 1 μ ASV for 24 h and then provided to Raji cells with three different ratios (10:1, 5:1, and 1:1) of effector sCAR19 T cells to target Raji cells. We pre-labeled Raji cells with calcein AM. After 4 h coculturing, we measured the levels of calcein AM that was released from Raji cells indicating cytolysis. Three control experiments using un-transduced T cells, CAR19 T cells, and sCAR19 T cells cultured in the absence of ASV were also set up as comparisons. As shown in Fig. 3a, both un-transduced T cells and sCAR19 T cells cultured in the absence of ASV exhibited very weak cytotoxic effects in killing Raji cells at all three tested effector-to-target ratios. However, both unswitchable CAR19 T cells and switchable sCAR19 T cells cultured in the presence of 1 μM ASV displayed strong cytotoxic effects in killing Raji cells and this killing effect was positively correlated to the effector-to-target ratio. At a 10:1 ratio, CAR19 T cells led to cytolysis of around 80% Raji cells in comparison to around 60% Raji cell cytolysis caused by sCAR19 T cells cultured in the presence of ASV. The slightly lower cytotoxic effect from sCAR19 T cells cultured in the presence of ASV is expected since residual HCV-NS3 activity at 1 *μ*M ASV will lead to a low level of sCAR19 cleavage. Also, the HCV-NS3 structure may influence the cytotoxicity.

**Fig. 3:**
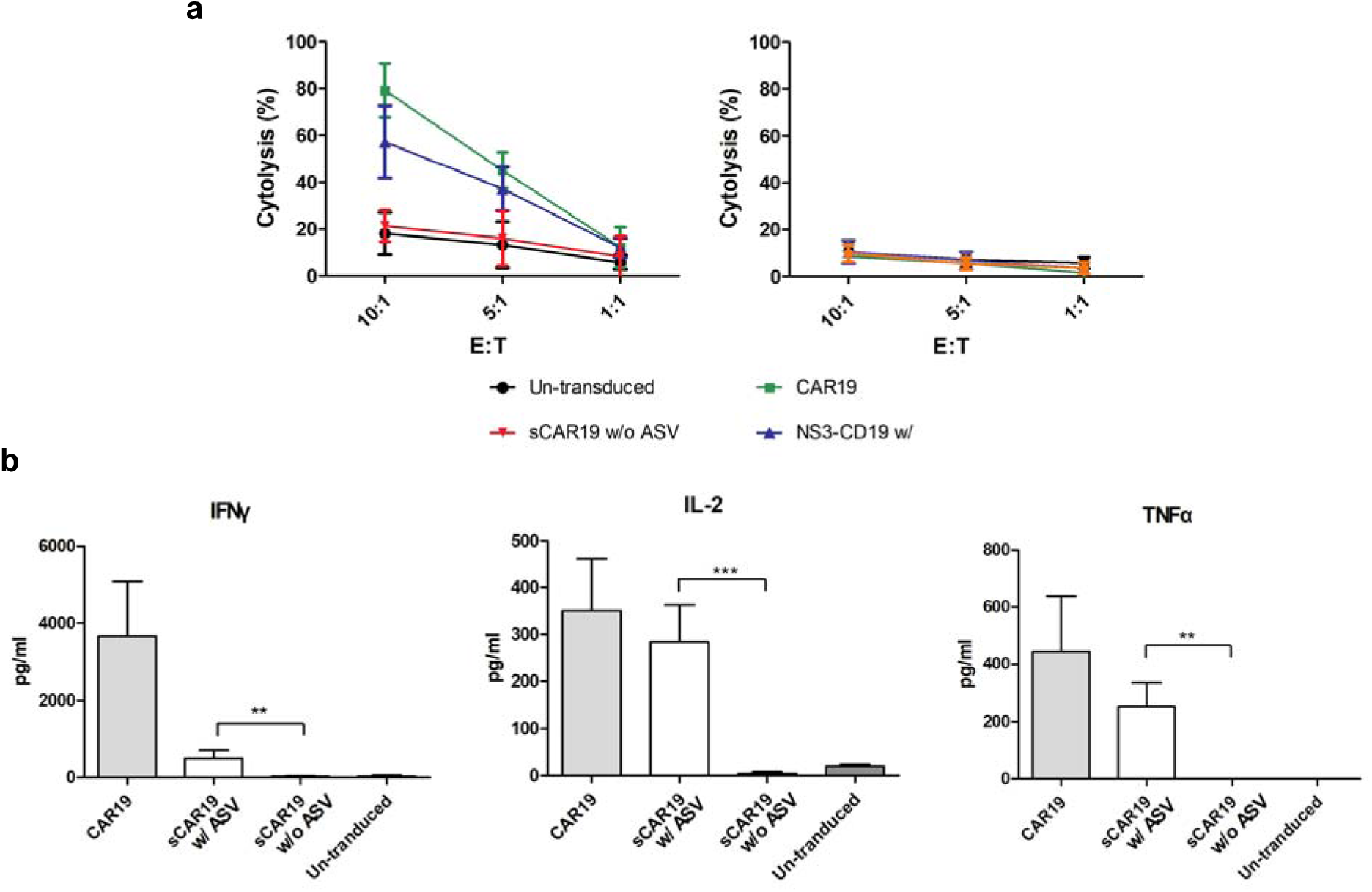
**sCAR19 T cells exhibit cytotoxicity *in vitro*.** (a) Cytotoxicity of un-transduced T cells, CAR19 T cells and sCAR19 T cells cultured in two conditions, one with the DMSO vehicle and the other with 1 μM ASV toward CD19^+^ Raji (the left graph) and CD19^-^ K562 cells (the right graph). Both Raji and K562 cells were labeled with calcein-AM before they were cocultured with four groups of T cells with ratios of effector to target tumor cells (E:T) as indicated in the figures for 4 h. Lysis of target cells was analyzed by detecting released calcein-AM in media. Data are representative of three independent experiments and normalized against total lysis of calcein-AM-labeled Raji and K562 cells. (b) The release of cytokines including IFN-γ, IL-2 and TNFα from un-transduced T cells, CAR19 T cells and sCAR19 T cells cultured in two conditions, one with the DMSO vehicle and the other with 1 μM ASV when they were cocultured with Raji cells with a E:T ratio as 1:1 for 24 h. Cytokine levels were detected using ELISA.

To study the influence of the recurring chemogenetic switch on T cell degranulation, an immune reaction process in response to a detected antigen, we cocultured sCAR19 T and Raji cells with and without 1 μM ASV for 4 h and then analyzed the expression of CD107a, a T cell degranulation marker on the T cell surface. Experiments with un-transduced and CAR19 T cells were set up as negative and positive controls. As a negative control, un-transduced T cells showed about 3% detectable labeling with a phycoerythrin (PE)-conjugated antibody for CD107a. On the contrary, CAR19 T cells as a positive control exhibited about 32% detectable labeling with the CD107a antibody. For sCAR19 T cells, about 24% and 5% were CD107a- positive for culturing conditions with and without 1 μM ASV respectively (Extended Data Fig. 7). These two degranulation levels that were similar to positive and negative controls respectively corresponded to active and inactive T cell antigen-response states pretty well. Replacing Raji cells (CD19^+^) with K562 (CD19^-^) cells led to no T cell degranulation for sCAR19 T cells that were cultured in the presence of 1 μM ASV. To see whether sCAR19 T cells display ASV-regulated long term anti-tumor effects at a low effect-to-target cell ratio, we pre-labeled Raji cells with carboxyfluorescein succinimidyl ester (CFSE) and then cocultured it together with sCAR19 T cells at a 1:10 effector-to-target cell ratio with and without 1 μM ASV for 3 days. Resulted cells were analyzed to calculate proportions of CFSE^+^ Raji cells. Similar experiments were set up with un-transduced and CAR19 T cells as negative and positive controls respectively. After 3 days of coculturing, proportions of CFSE^+^ Raji cells in the un-transduced, CAR19, ASV- absent sCAR19, and ASV-present sCAR19 T group were around 56%, 18%, 53%, and 15% respectively (Extend Data Fig. 8). Correspondingly, T cell proportions in the four groups were 42%, 81%, 44%, and 84% respectively. This result showed convincingly that ASV-treated sCAR19 T cells behaved similarly as CAR19 T cells in inducing a long-term anti-tumor effect and withdrawing ASV drove sCAR19 T cells completely to an inert state similar to un-transduced T cells. Collectively, data in this section demonstrate that the recurring chemogenetic switch can shift the sCAR19 T cells between active and inactive T cell states for implementing controllable long-term cytotoxicity effects in eliminating tumor cells *in vitro*.

### ASV-controlled anti-tumor effect of switchable CAR-T cells in mice

Given the clear ASV- regulated anti-tumor effect of sCAR19 T cells *in vitro*, we advanced our study in an animal model to evaluate the said effect *in vivo* according to a chart presented in Extend Data Fig. 9. 5×10^5^ Raji-Luc cells that stably express luciferase for bioluminescent imaging were infused into 40 mice via intravenous tail injection to induce a lymphoma tumor phenotype. We raised these tumor-infused mice for 7 days and then separated mice to three groups for intravenous infusion with un-transduced, CAR19, and sCAR19 T cells. We further separate mice in the sCAR19 T group into three subgroups for gavage feeding with three different daily doses of ASV as 0 (a 9:1 PEG400:ethanol vehicle), 2, and 15 mg/kg. To evaluate whether ASV alone provides tumor eradication effect, we gavage fed mice infused with un-transduced T cells with a daily ASV dose as 15 mg/kg. To assess tumor engraftment and antitumor activity of different T cells, we evaluated tumor growth in each mouse by detecting luciferase-catalyzed bioluminescence in its body in different days and using anti-human CD3 and CD19 antibodies to detect human T and Raji cells respectively once a week. In the CAR19 T group that was used as a positive control, 3 out 4 mice survived beyond day 28. Tumor cell prevalence reached its peak around day 16 after the Raji cell injection and decreased significantly after day 20 indicating a strong anti-tumor effect from the infused CAR19 T cells (Fig. 4a). In the absence of ASV, mice infused with sCAR19 T cells all reached demise at day 20. When gavage feeding with 2 mg/kg ASV daily, the mortality of sCAR19 T-infused mice was significantly decreased and 2 mice survived beyond day 20. When the fed ASV was improved to 15 mg/kg daily, all four mice survived beyond day 20 and two survived beyond day 28 with tumor cells significantly lower than at day 20. Mice in the un-transduced T cell group that was also fed with a daily dose of ASV as 15 mg/kg all reached demise at day 20 indicating that ASV alone does not provide significant tumor killing effects. The analysis of tumor cell and T cell percentages in the blood from mice infused with different T cells indicated that in the absence of ASV, mice in the sCAR19 T group had tumor Raji and human T cells at levels similar to mice treated with un-transduced mock T cells (Fig. 4c and Extend Data Fig. 10). This indicated that both tumor Raji and human T cells proliferated at similar rates in two groups of mice and both un-transduced mock T and ASV-untreated sCAR19 T cells did not implement tumor killing effects. The tumor killing effect was significantly improved when ASV was gavage-fed to Raji-grafted mice at a 2 mg/kg daily dose and reached to a level close to the positive-control CAR19 T cells when the ASV dose reached 15 mg/kg (Fig. 4b and Extended Data Fig. 10). The activation of sCAR19 T cell proliferation was also observed when ASV was injected to Raji-grafted mice and reached to a level close to that from the positive control CAR19 T cells when the ASV dose reached 15 mg/kg (Fig. 4c and Extended Data Fig. 10). The analysis of lentiviral copy numbers per ug of DNA isolated from the blood of mice in different groups also indicated strong ASV-induced sCAR19 T proliferation in mice. Collectively, our results support strongly a robust ASV-controlled anti-tumor effect of sCAR19 T cells *in vivo*.

**Fig. 4:**
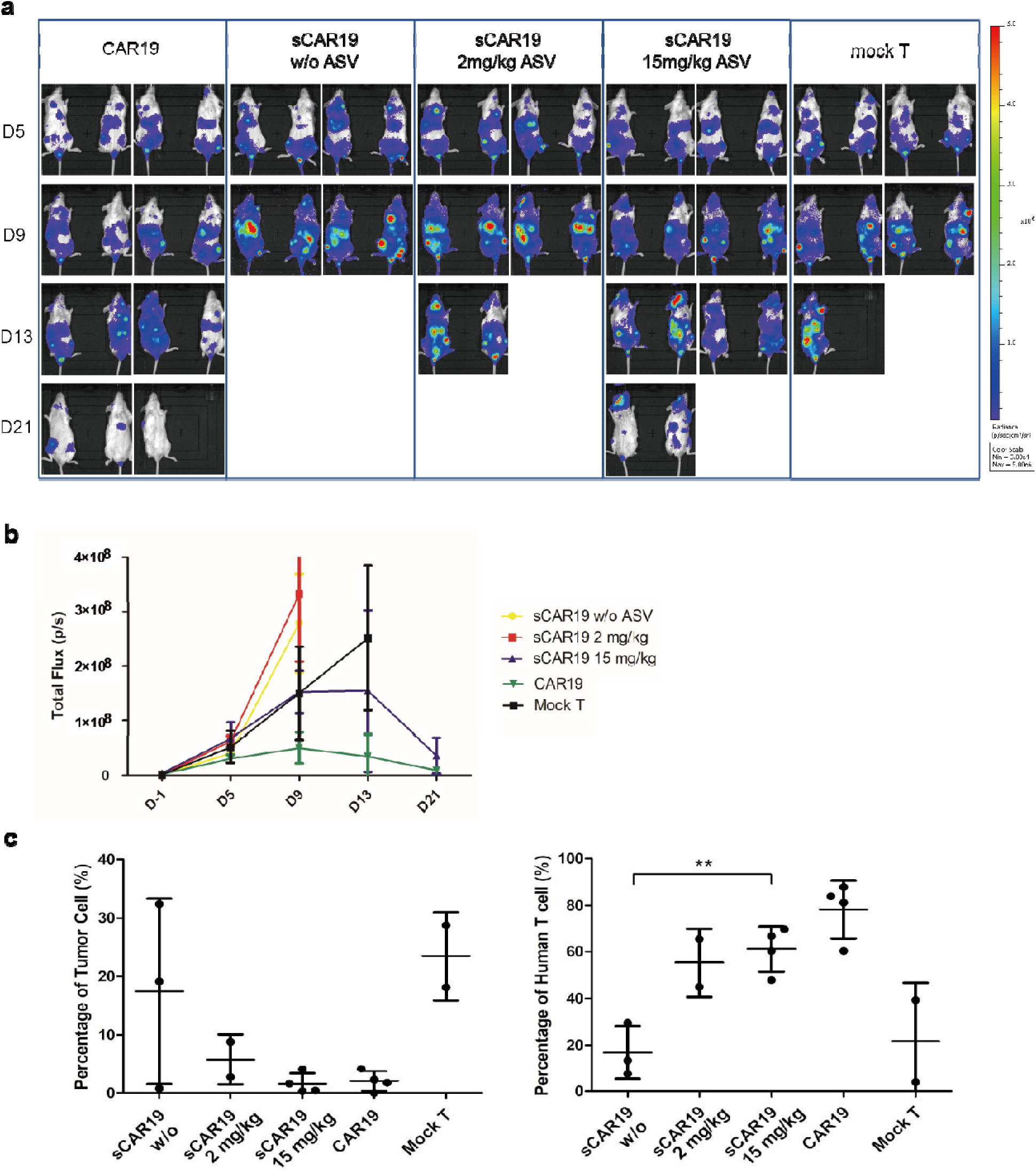
**Switchable CAR-T cells in combination with ASV are effective in eliminating human CD19^+^ tumor cells in mice. (a) Tumor growth in mice that were monitored by bioluminescent imaging. Mice were infused with Raji-Luc cells (5×10^5^ cells per mouse) at day 0 and then treated with un-transduced human T cells, CAR19 T cells and sCAR19 T cells (5×10^6^ cells per mouse) at day 7. Mice treated with un-transduced T cells were fed daily with 15 mg/kg ASV, mice treated with CAR19 T cells were fed daily with a vehicle (PEG400:ethanol as 9:1) and mice treated with sCAR19 T cells were fed daily with the vehicle, 2 mg/kg ASV and 15 mg/kg ASV. To image tumor growth, mice were anesthetized and then injected with D-luciferin to undergo whole body bioluminescent imaging.** (b) Tumor cell growth and elimination indicated by luciferase-catalyzed bioluminescence. Whole body bioluminescence for each survived mouse that was detected from the ventral side was used for the calculation. Y axis indicates the average bioluminescent signals for each survived mouse. (c) Raji and human T cell counts in blood from mice with different treatments at day 18. The percentages were determined with respect to all leukocytes in collected blood.

## Conclusions

Unlike small molecules with well-defined pharmacokinetic/pharmacodynamic features, the unpredictability of CAR-T cell therapeutics faces unique treatment challenges regarding the dose control. The ability to regulate the activity and survival of these live drugs has dual benefits of promoting efficacy without abandoning safety^33, 34^. By introducing a recurring chemogenetic switch into the CAR that can be regulated through a small molecule, the problem turns into control of the dosage of the small molecule, which in the current work is a well-studied FDA approved HCV-NS3 inhibitor ASV. Using ASV to regulate activity of HCV-NS3 embedded in sCAR19, we show that full-length sCAR19 was displayed on its host CAR-T cell surface in an ASV-dose dependent manner. More importantly, we found that the removal of ASV resulted in gradual elimination of the CAR, and as fast as in 24 h the CAR level decreased to an insignificant level. Most biological characterization results prove that CAR-T cells with this recurring chemogenetic switch behave similarly to standard CAR-T cells and ASV regulates, with a dose dependent manner, the antitumor effect of switchable CAR-T cells both *in vitro* at the cellular level and *in vivo* in mice. Due to its recurring control nature, we believe this novel chemogenetic CAR switch will find critical applications in CAR-T cell therapy in mitigating serious side effects such as cytokine release syndromes and T cell exhaustion. Since ASV is an FDA-approved medication and sCAR19 in the current work was derived from an approved CAR-T therapeutic, the combined use of ASV and sCAR19 as a therapeutic can be potentially advanced quickly to undergo clinical investigations in the treatment of hematopoietic tumors such as B-cell lymphoblastic leukemia.

## Online content

Any methods, additional references, Nature Research reporting summaries, source data, extended data, supplementary information, acknowledgements, peer review information; details of author contribution and competing interests; statements of data and code availability are available.

## Data availability

The authors declare that the main data supporting the findings of this study are available within the article and its Supplementary Information files. The expression plasmids are available from W.R.L.

## Acknowledgements

The project was sponsored by IASO USA Inc. Z.Z.G. was partially supported by a Welch Foundation grant A-1715. The mouse study was carried out in the animal facility of Comparative Medicine Program at Texas A&M University.

## Author contributions

J.Z. and W.R.L. designed the research. W.C., Z.Z.G. and Q.P. performed the research and collected data. S.G., S.X., J.Z. and W.R.L. analyzed and interpreted the data. W.C., J.Z. and W.R.L. wrote the manuscript. All authors critically revised the manuscript.

## Methods

### Plasmid construction

We ordered two DNA fragments, one coding anti-human CD19 scFv and the other coding CAR domains including the hinge, transmembrane domain and cytoplasmic regions of human CD28, 4-1BB, and CD3ζ from IDT DNA Inc. Sequence information of the two fragments can be found in Supplementary Table 1. We did overlap PCR to ligate the two DNA fragments to make the full-length CAR19 DNA using primers CAR19-F and CAR19-R (primer sequences can be found in Supplementary Table 2), digested the amplified DNA using restriction enzymes EcoRI and MluI, and ligated the digested DNA into the lentiviral plasmid pLVX-EF1α-IRES-puro between sites EcoRI and MluI to afford pLVX-EF1α CAR19. The plasmid map of pLVX-EF1αCAR19 is shown in Supplementary Fig. 1. We acquired pLVX-EF1α-IRES-puro from Takara Bio Inc. Its digestion by enzymes EcoRI and MluI removed the coding regions for IRES and the puromycin resistance gene. To add the HCV-NS3 coding sequence into the CAR region of pLVX-EF1α AR19, we ordered its DNA fragment from IDT DNA Inc., amplified it using primers NS3-F and NS3-R (Supplementary Table 2), digested it with SpeI and XbaI, and then cloned the digested DNA into pLVX-EF1α CAR19 between SpeI and XbaI to afford plasmid pLVX-EF1α CAR19. In order to build a tumor cell line that stably expresses luciferase, we constructed another plasmid pLVX-puro-Luc. We ordered a DNA fragment coding firefly luciferase (its sequence in Supplementary Table 1), amplified it using primers Luc-F and Luc-R (Supplementary Table 2), digested it with EcoRI and XbaI, and then cloned into pLVX-EF1α IRES-puro between restriction sites EcoRI and XbaI. We confirmed all constructed plasmids using DNA sequencing services provided by Eton Bioscience Inc.

### Cell line culture

We purchased K562, Raji and HEK 293T/17 from American Type Culture Collection. We maintained K562 and Raji cell lines using RPMI 1640 medium with 10% fetal bovine serum (FBS). We cultured HEK 293T/17 with high glucose DMEM medium with 10% FBS. Both media and FBS were acquired from Gibco Inc. All cells were cultured at 37 °C with 5% CO_2_. Cells at logarithmic growth phase were used for following experiments.

### Lentivirus packaging and concentration

We prepared all plasmids for lentivirus packaging using the EndoFree Plasmid Midi Kits from Omega Bio-tek Inc. according to the manufacturer’s protocol. For packaging unswitchable CAR19 lentivirus particles, we grew HEK293T/17 cells in ThermoFisher 10 cm dishes to 70-80% confluency and then co-transfected cells with three plasmids pLVX-EF1α CAR19, psPAX2 and PMD2.G using polyethyleneimine (Polysciences, Inc.) as described previously^1^. We acquired psPAX2 encoding lentiviral packaging proteins and PMD2.G encoding a lentiviral envelop protein from Addgene (plasmid no. 12260 and 12259 respectively). We grew the transfected cells for 2 days and collected supernatants (10 mL) to isolate viral particles. Additional medium (10 mL) was provided to transfected cells for growing one more day and subsequently we collected supernatants. We centrifuged the collected supernatants at 4000× g at 4 °C to remove cell debris, filtrated the residual supernatants through a 0.45 μm membrane, and then centrifuge at 30,000× g at 4 °C to precipitate viral particles. We then removed the supernatants and resuspended the lentiviral pellets in DMEM medium (200 μL for a total of 200 mL collected supernatant volume). We aliquoted resuspended lentiviral particles as 200 μL/each and stored them at -80 °C. We detected p24 on lentiviral particles using Lenti-X GoStix Plus Kits provided by Takara Inc. to confirm successful production of lentivirus and titered collected lentiviral particle-containing solutions using 293T cells following standard protocols and detecting CAR19 expression on infected 293T cells using Alexa Fluor 647-labeled rabbit anti-mouse F(ab)^2^. To package switchable CAR lentiviral particles, we followed the exact same protocol except replacing plasmid pLVX-EF1α CAR19 with plasmid pLVX-EF1α sCAR19. We also produced lentiviral particles for transduce Raji cells to make stable Raji cells expressing luciferase. We followed the same protocol by replacing plasmid pLVX-EF1α CAR19 with plasmid pLVX-puro-Luc.

### T cell isolation, transduction and culture

We purchased leukocyte products from the Gulf Coast Regional Blood Center. In order to isolate T cells, we typically added 20 mL leukocyte products on top of 20 mL Ficoll-Paque solution for density gradient centrifugation from GE Healthcare in a 50 mL Falcon tube and then centrifuged at 800× g at room temperature for 20 min to separate different leukocyte cells. We then collected peripheral blood mononuclear cells (PBMCs) and washed collected cells with a 40 mL sorting buffer containing 1× phosphate buffer saline (PBS) and 2 mM EDTA at pH 7.2 twice and spun down the cells. To isolate T cells from PBMCs, we suspended pelleted PBMCs in the sorting buffer (80 μL for 10^7^ cells) with CD3 magnetic microbeads (20 μL for 10^7^ cells) from Miltenyi Biotec for 15 min at 4 °C according to the capacity of the microbeads. Cell numbers were determined using the standard cell counting approach. We centrifugated the mixture to remove the supernatants and then washed the microbead pellets once with the sorting buffer. We suspended microbead pellets in additional sorting buffer (500 μL for 10^8^ cells) and then loaded the suspended microbeads to a LS column (Miltenyi Biotec). We washed the column that was attached to a magnetic MidiMACS^TM^ separator from Miltenyi Biotec for retaining CD3 magnetic microbeads with the sorting buffer three times to remove CD3-negative cells and then transferred the beads bound with T cells to a 15 mL falcon tube. Cells were washed with the sorting buffer once and suspended in ThermoFisher CTS™ OpTmizer™ medium (Gibco) supplemented with 2 mM L-glutamine (Gibco) in the presence of 200 IU/ml human IL-2 (PeproTech). We then stimulated T cells with Dynabeads™ Human T-Activator CD3/CD28 from ThermoFisher overnight with a ratio of beads to cells as 1:3. To transduce T cells, we incubated them with suspended lentiviral solutions (MOI 3-5) for 24 h with 4 µg/mL polybrene (Sigma-Aldrich) before we replaced the medium with refresh culture medium. T cells were maintained at a density of 0.5-2×10^6^/mL at all times.

### CAR detection on transduced T cells

For anti-human CD19 scFv detection, we labeled transduced T cells with Alexa Fluor 647- labeled rabbit anti-mouse F(ab)^2^ antibody from Jackson ImmunoResearch Laboratories Inc. or FITC-labeled human CD19 from AcroBiosystems following the manufacture’s protocols. All antibodies and their providers that we used in the current study are listed in Supplementary Table 3. We sorted labeled cells using a CytoFLEX flow cytometer (Beckman Coulter). CAR-positive cells were gated using un-transduced cells as controls. By subtracting the percentage of positive cells in control samples from the percentage of positive cells in CAR19 transduced samples, the background was eliminated. For sCAR19 T cells, we cultured them in various concentrations of ASV for different times before labeling and sorting.

### T cell proliferation, subset and apoptosis

To test the proliferation of CAR-T cells, we used carboxyfluorescein diacetate succinimidyl ester (CFSE, ThermoFisher) to label the cells according to the manufacturer’s instruction. The apoptosis assay was conducted using an Annexin V Apoptosis Detection Kit (BD Biosciences) according to the manufacturer’s instruction. To analyze T cell subsets, we stained T cells with three different color-labeled anti-CD4 and anti-CD8 antibodies from BioLegend Inc. and sorted them using the CytoFLEX flow cytometer. Flow cytometry data analysis was performed using FlowJo software from TreeStar.

### The characterization of T cell cytotoxicity by detecting Calcein-AM release

The Calcein-AM release assay was conducted according to a previously reported method^3^. Briefly, the positive target cells (Raji) and negative target cells (K562) were labeled with Calcein-AM (Biolegend) and then cocultured with effector cells (CAR19 and sCAR19 cells) in 96-well plates at different ratios with PBS + 5% FBS. Spontaneous release wells were set as cocultured target cells and PBS + 5% FBS, and maximum release wells were target cells and lysis solution. After 4 h incubation at 37 LJ, the plate was centrifuged at 500× g for 10 min and the supernatants were transferred to another 96-well plate. The fluorescence value of each well (R) was measured with the microplate reader, and the tumor-killing efficiency was calculated by the following formula: Lysis% = (R_experimental well_ - R_spontaneous release_)/(R_maximum release_ - R_spontaneous release_) × 100%.

### The characterization of T cell cytotoxicity by detecting CD107a expression on the T cell surface

To characterize CD107a expression on T cells, they were cultured together with Raji and K562 cells at 37 °C for 4 h in 24-well plates. For each culture, we added 20 μL original PE-conjugated anti-CD107a solution purchased from ThermoFisher and 1 μL of Golgi Stop (monesin) from BD Biosciences to 500 μL culture medium. For sCAR19 T cells, various concentrations of ASV were provided. We then stained T cells with FITC-conjugated anti-CD3 and sorted them using the CytoFLEX flow cytometer. We analyzed Collected data using FlowJo.

### The characterization of cytokines including IFN-**γ**, TNF-**α**, and IL-2 released to the growth media using ELISAs

We co-cultured 5×10^5^ effector T cells with the same number of target cells, both Raji and K562 cells, in 24-well plates. The plates were incubated at 37 °C for 24 h. After that, we transferred supernatants and characterized their IFN-γ, TNF-α, and IL-2 using ELISA kits provided by ThermoFisher. The characterizations followed the manufacturer’s protocols. When two CARs were compared, cytokine release was normalized for CAR expression by dividing the cytokine levels by the fraction of CAR expression. For sCAR19 T cells, various concentrations of ASV were provided to the growth media.

### The establishment of Raji-Luc cells

To establish Raji-Luc cells that stably express luciferase, we used lentivirus particles that were packaged from the use of pLVX-puro-Luc to infect Raji cells and cultured the infected cells in the presence of 2 μg/mL puromycin from Gibco Inc. for a week to select stable cells. We verified luciferase expression in selected stable cells after providing 150 ug/ml D-luciferin to the growth media and detecting bioluminescence using an IVIS In Vivo Imaging System (PerkinElmer).

### Murine lymphoma experiments

Animal studies were approved by the Texas A&M’s Institutional Animal Care and Use Committees. We purchased 5-week-old female NSG mice (NOD.Cg-Prkdcscid Il2rgtm1Wjl/SzJ) from JAX laboratory. We built a CD19 positive tumor model in these mice by engrafting 5×10^5^ Raji-Luc cells through tail vein injection. Tumors were allowed to grow for 6 days, and then we infused the mice with 5×10^6^ un-transduced, CAR19 or sCAR19 T cells. ASV at two doses as 2 and 15 mg/kg or a vehicle (PEG400 to ethanol as 9:1) was given by oral gavage once a day after the sCAR19 T cell infusion. For the un-transduced T cell control group of mice, 15 mg/kg ASV was given by oral gavage once a day and for the positive CAR19 T control group, the vehicle was given by oral gavage once a day. We evaluated *in vivo* tumor growth through bioluminescent imaging using the IVIS In Vivo Imaging System twice a week. Before bioluminescent imaging, we anesthetized mice with isoflurane and gave them by intraperitoneal injection 150 mg/kg D-Luciferin potassium salt from MedChemExpress. Bioluminescent images of whole mice were taken 10 minutes after D-luciferin injection and these mice were recorded. Once in two weeks, we collected blood cells from these mice, stained them with FITC-conjugated anti-human CD3 for the detection of human T cells and PE-conjugated anti-human CD19 for the detection of survival Raji-Luc cells, and then sorted them using the CytoFLEX flow cytometer. Percentages of human T cells and tumor cells in all sorted leukocytes were then determined. We also extracted DNA from collected mouse blood and quantified their lentiviral levels by running real-time PCR using primers Lenti-F and Lenti-S. Lentiviral DNA copies per μg overall extracted DNA were then determined. Survived mice were counted every day and the mortality was calculated.

### Statistical analysis

The results are presented as the means ± the standard deviations. Student’s t-tests were used for data comparisons as indicated in the figure legends. GraphPad Prism 7 were used for all statistical analyses. The number of samples in each experiment is indicated in the figure legends. P < 0.05 was considered statistically significant.

## Additional information

### Extended data are available for this paper

**Extended Data Fig. 1:**
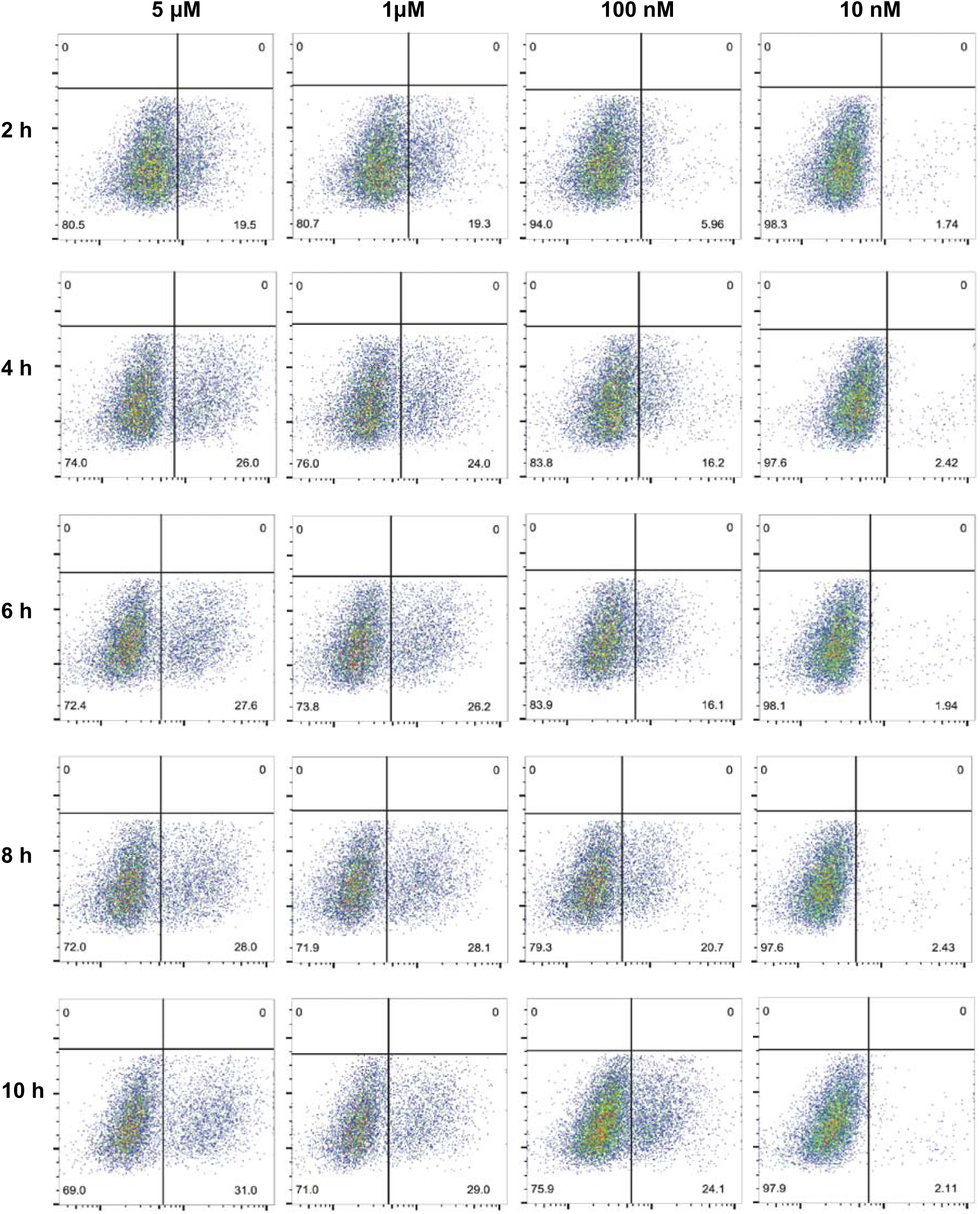
**Flow cytometry results of full-length sCAR19 display on T cells at different timepoints and ASV concentration. Alexa Fluor 647-anti-mouse F(ab)2 antibody was used for the detection of full-length sCAR19 display. The bottom right section of each dot plot shows cells with expressed full-length sCAR19.**

**Extended Data Fig. 2:**
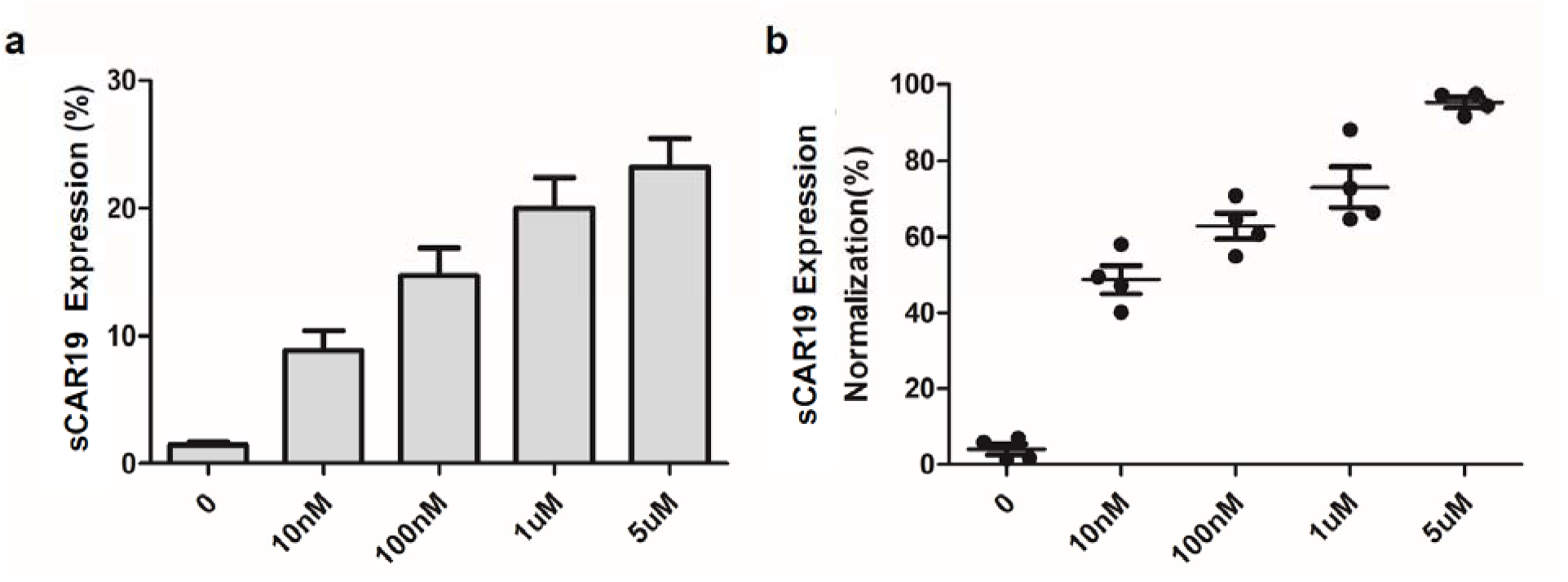
**Dose-dependent full-length sCAR19 display on T cells. a. Full-length sCAR19 levels on T cells in the presence of 0, 10 nM, 100 nM, 1 μM and 5 μM ASV at the 10 h time point. b. The displayed full-length sCAR19 on T cells after normalization in the presence of 5μM ASV at the 24 h time point.**

**Extended Data Fig. 3:**
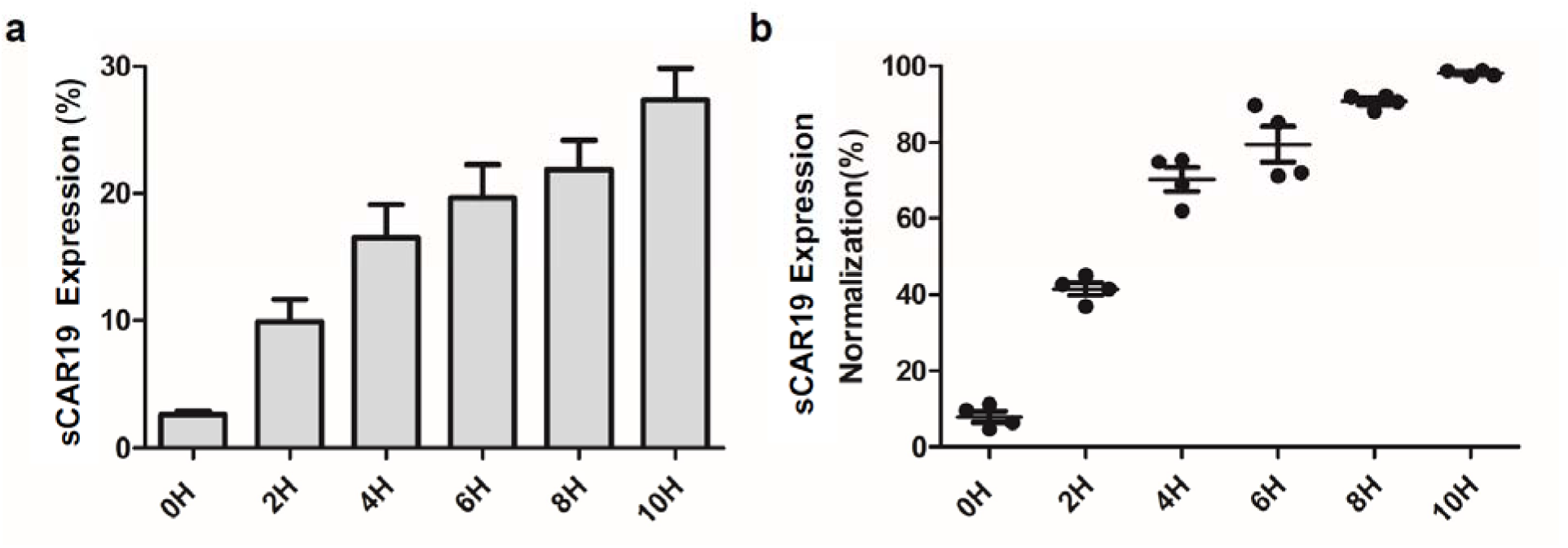
**Time-dependent full-length sCAR19 display on the T cell surface in the presence of ASV. a. Full-length sCAR19 levels on the T cell surface at 0, 2, 4, 6, 8 and 10 h time points after the addition of 5 μM ASV. b. Displayed sCAR19 levels after normalization in the presence of 5 μM ASV at different time points.**

**Extended Data Fig. 4:**
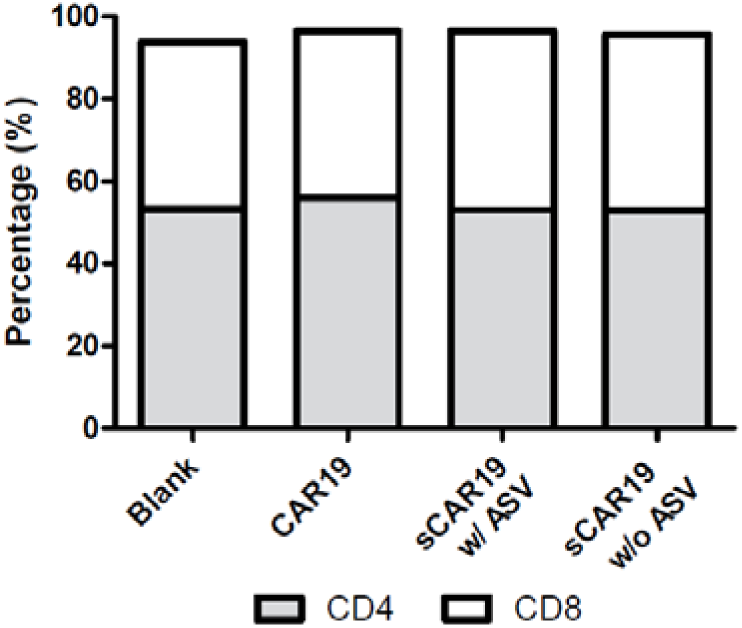
**Subsets of sCAR19 T cells. Four groups T cells were cultured with or without 5 μM ASV for 72 h. CD4 and CD8 percentage were analyzed by flow cytometry.**

**Extended Data Fig. 5:**
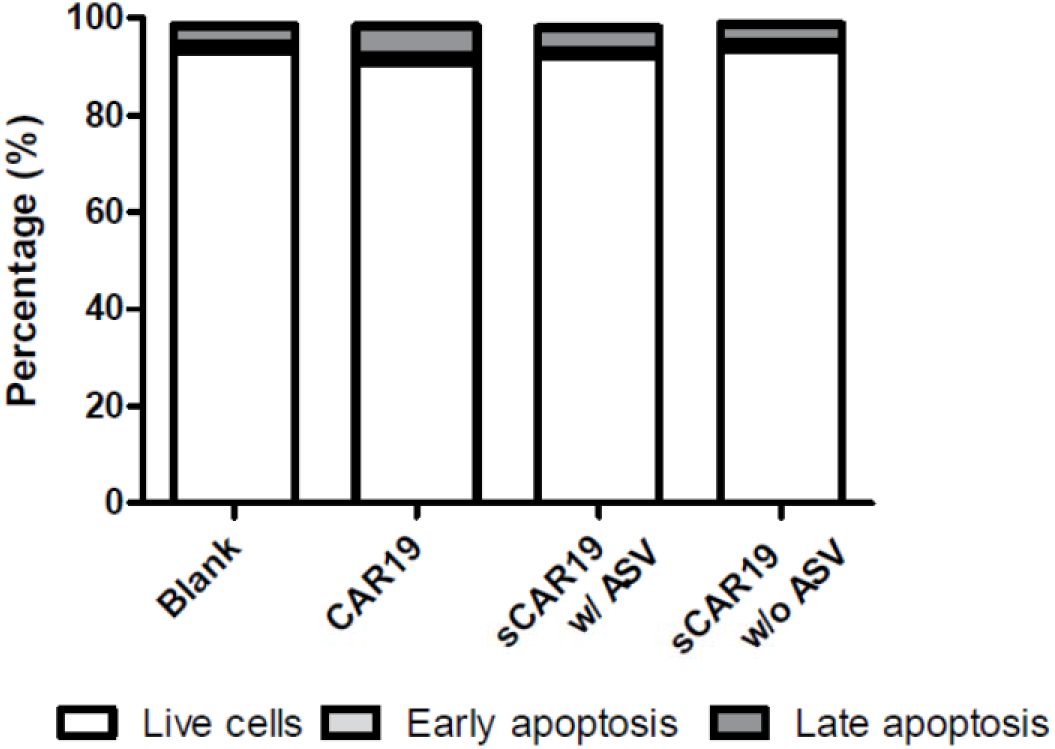
**Apoptosis of sCAR19 T cells in the presence of 5 μM ASV. Apoptosis was tested after 3 days of ASV incubation. The apoptotic rate of each group showed no significant difference (P >0.05).**

**Extended Data Fig. 6:**
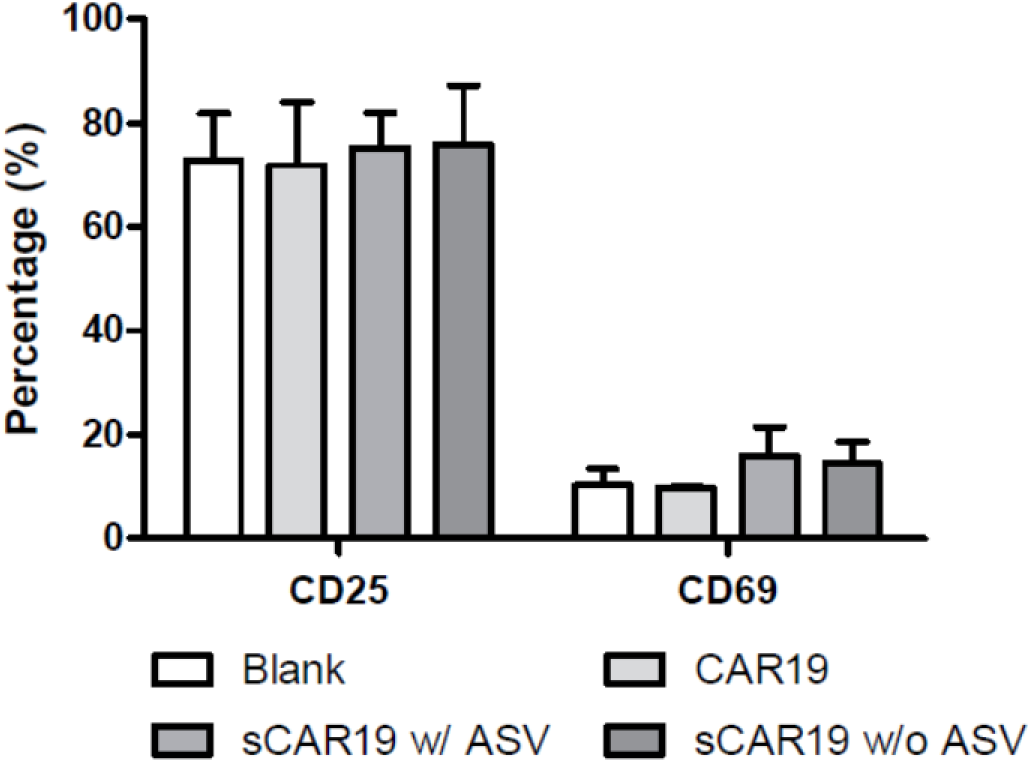
**Activation of sCAR19 T cells. CD25 and CD69 were analyzed by flow cytometry after three days culture. The expression of CD25 and CD69 showed no significant difference in four groups (P > 0.05). ASV was provided as 5 μM.**

**Extended Data Fig. 7:**
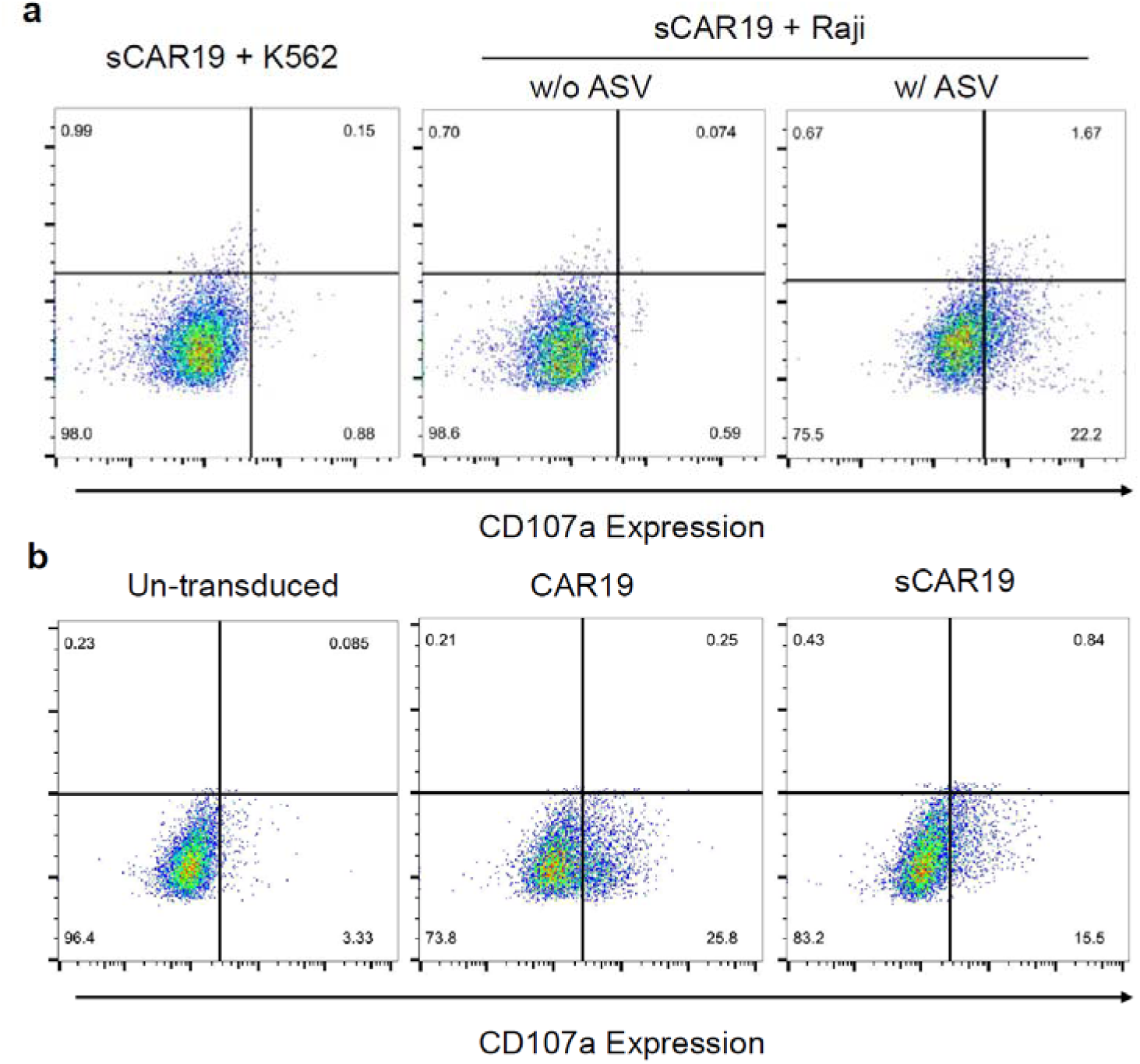
Degranulation analysis of sCAR19 T cells by the detection of CD107a expression. sCAR19 T and target cells were cocultured with and without 1 μM ASV for 4 h and analyzed by flow cytometry. The plots are gated on CD3^+^ portions. a. sCAR19 cocultured with K562 (CD19^-^) and Raji (CD19^+^) with or without 1 μM ASV. For K562 cells, only data in the presence of 1 μM ASV are shown. b. Un-transduced, CAR19 and sCAR19 T cells cocultured with Raji with 1μM ASV.

**Extended Data Fig. 8:**
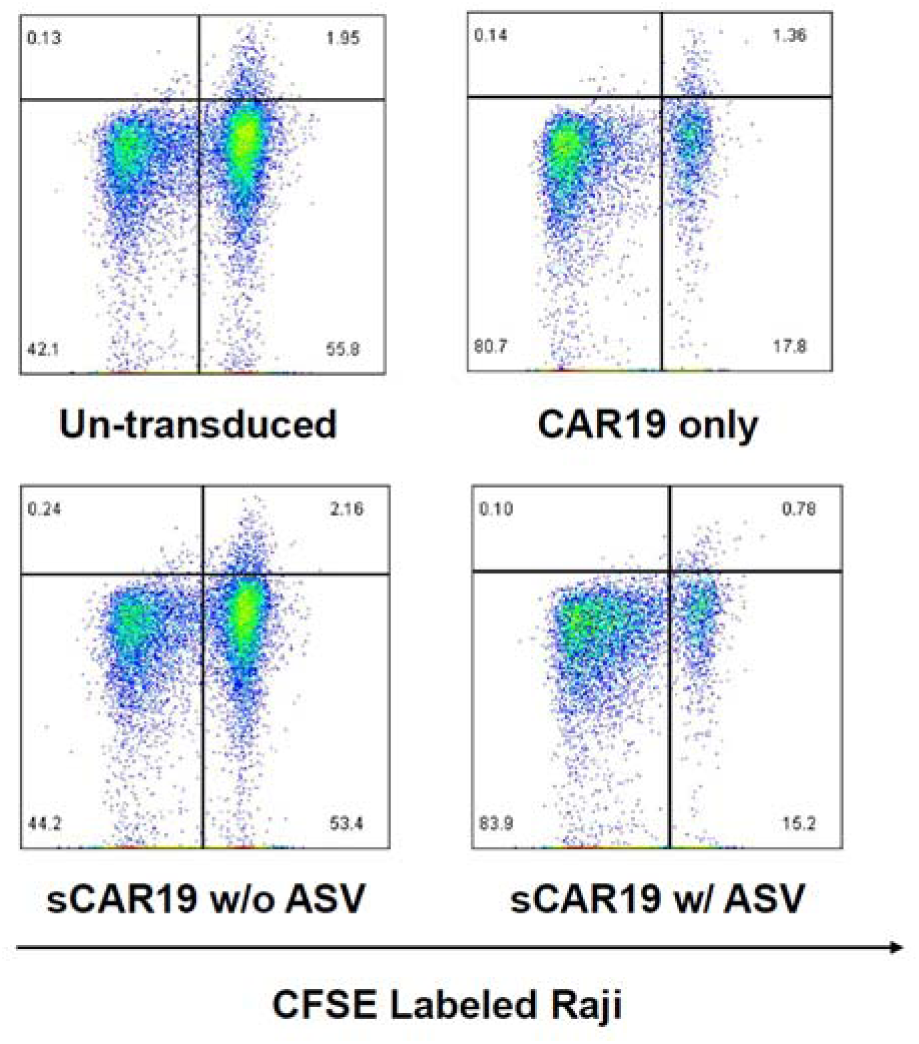
**Long-term antitumor effects of sCAR19 T cells in the presence of 1 μM ASV. CFSE labeled Raji cells were cocultured with CAR19 or sCAR19 at low ratio of effector to target cells (E: T = 1: 10). After 72 hours of coculturing, the proportion of CFSE^+^ Raji cells was detected by flow cytometry.**

**Extended Data Fig. 9:**
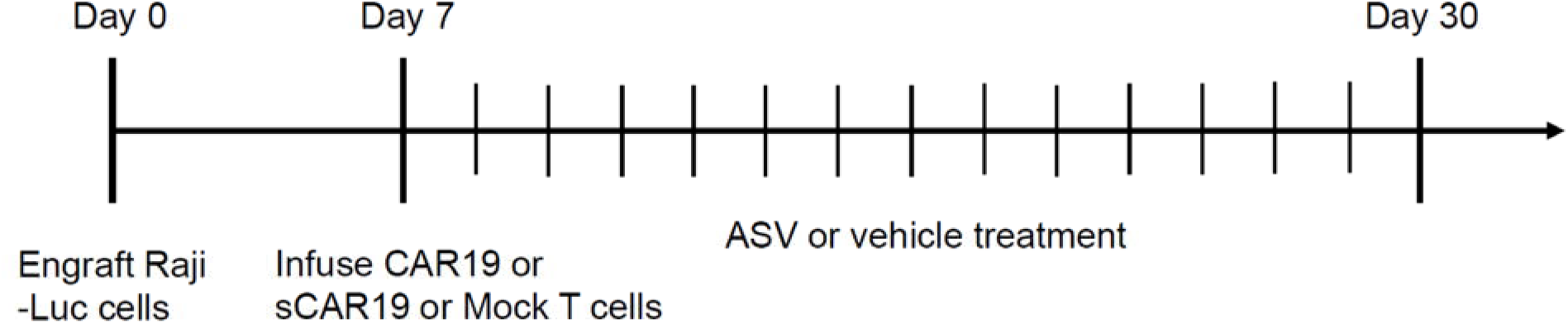
**Flowchart of the mouse study. Raji-Luci cells were engrafted on Day 0, then CAR19, sCAR19, or Mock T cells were infused according to the group. ASV or vehicle was given once per day since T cell infusion until Day 30.**

**Extended Data Fig. 10:**
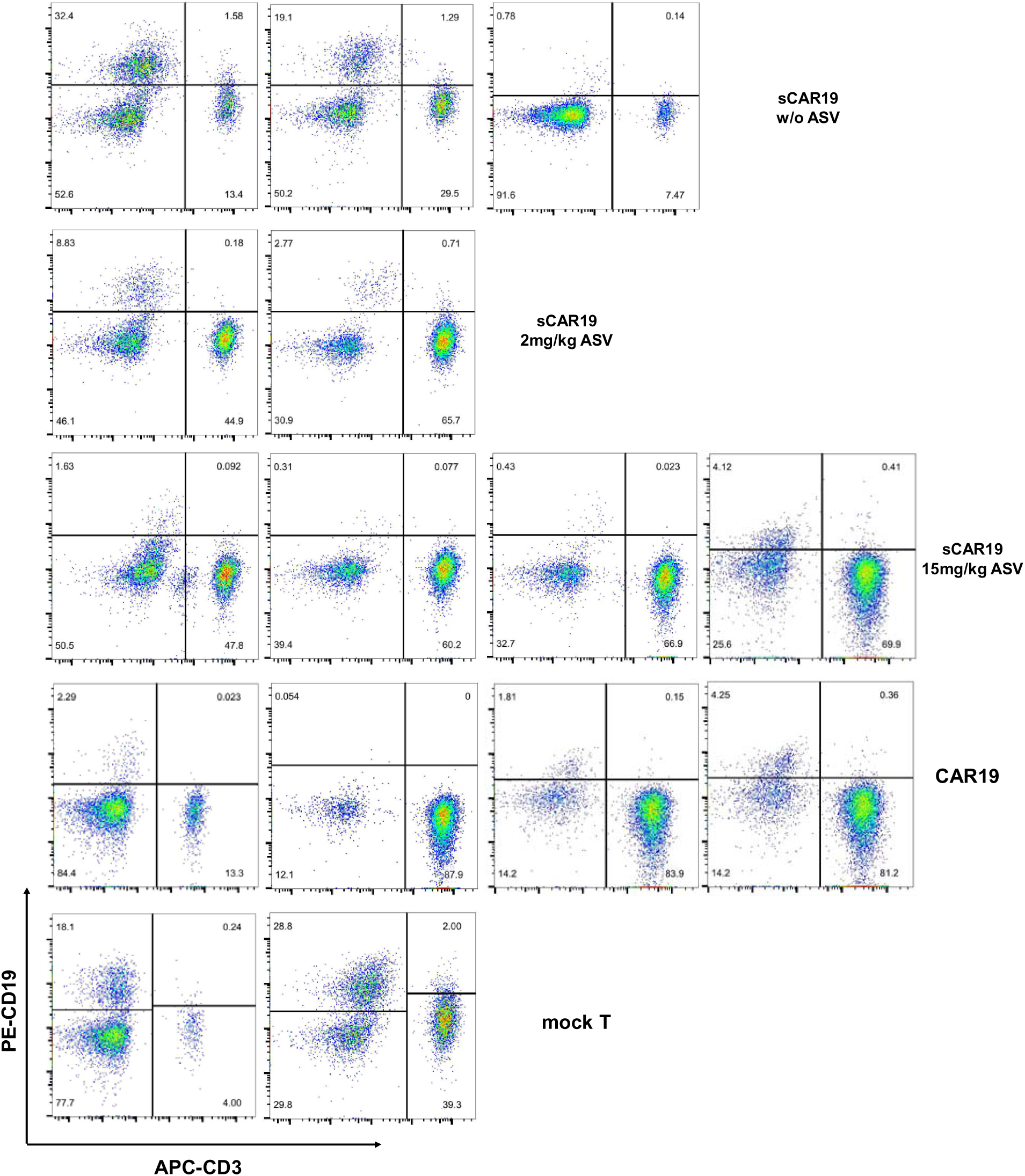
**Flow cytometry results of CD3 and CD19 detection of blood in all individual survived mice in different groups in Day 18. Each dot plot represents data for each particular mouse. APC-CD3 and PE-CD19 were used to differentiate Raji tumor cells and human T cells.**

## References

1. Maude, S.L. et al. Chimeric antigen receptor T cells for sustained remissions in leukemia. New England Journal of Medicine 371, 1507–1517 (2014).

2. Lee, D.W. et al. T cells expressing CD19 chimeric antigen receptors for acute lymphoblastic leukaemia in children and young adults: a phase 1 dose-escalation trial. The Lancet 385, 517–528 (2015).

3. Turtle, C.J. et al. CD19 CAR-T cells of defined CD4+:CD8+ composition in adult B cell ALL patients. The Journal of clinical investigation 126, 2123–2138 (2016).

4. Crump, M. et al. Outcomes in refractory diffuse large B-cell lymphoma: results from the international SCHOLAR-1 study. Blood 130, 1800–1808 (2017).

5. Neelapu, S.S. et al. Axicabtagene ciloleucel CAR T-cell therapy in refractory large B-cell lymphoma. New England Journal of Medicine 377, 2531–2544 (2017).

6. Turtle, C.J. et al. Immunotherapy of non-Hodgkin’s lymphoma with a defined ratio of CD8+ and CD4+ CD19-specific chimeric antigen receptor–modified T cells. Science translational medicine 8, 355ra116–355ra116 (2016).

7. Pan, J. et al. High efficacy and safety of low-dose CD19-directed CAR-T cell therapy in 51 refractory or relapsed B acute lymphoblastic leukemia patients. Leukemia 31, 2587–2593 (2017).

8. Boyiadzis, M.M. et al. Chimeric antigen receptor (CAR) T therapies for the treatment of hematologic malignancies: clinical perspective and significance. Journal for immunotherapy of cancer 6, 137–137 (2018).

9. Locke, F.L. et al. Phase 1 Results of ZUMA-1: A Multicenter Study of KTE-C19 Anti-CD19 CAR T Cell Therapy in Refractory Aggressive Lymphoma. Mol Ther 25, 285–295 (2017).

10. Majzner, R.G. & Mackall, C.L. Clinical lessons learned from the first leg of the CAR T cell journey. Nat Med 25, 1341–1355 (2019).

11. Wang, D. et al. A Phase I Study of a Novel Fully Human BCMA-Targeting CAR (CT103A) in Patients with Relapsed/Refractory Multiple Myeloma. Blood (2021).

12. Schuster, S.J. et al. Tisagenlecleucel in Adult Relapsed or Refractory Diffuse Large B-Cell Lymphoma. N Engl J Med 380, 45–56 (2019).

13. Lee, D.W. et al. Current concepts in the diagnosis and management of cytokine release syndrome. Blood 124, 188–195 (2014).

14. Shimabukuro-Vornhagen, A. et al. Cytokine release syndrome. J Immunother Cancer 6, 56 (2018).

15. Santomasso, B.D. et al. Clinical and Biological Correlates of Neurotoxicity Associated with CAR T-cell Therapy in Patients with B-cell Acute Lymphoblastic Leukemia. Cancer discovery 8, 958–971 (2018).

16. Wang, N. et al. Efficacy and safety of CAR19/22 T-cell cocktail therapy in patients with refractory/relapsed B-cell malignancies. Blood 135, 17–27 (2020).

17. Kasakovski, D., Xu, L. & Li, Y. T cell senescence and CAR-T cell exhaustion in hematological malignancies. J Hematol Oncol 11, 91 (2018).

18. Philip, B. et al. A highly compact epitope-based marker/suicide gene for easier and safer T-cell therapy. Blood 124, 1277–1287 (2014).

19. Gargett, T. & Brown, M.P. The inducible caspase-9 suicide gene system as a “safety switch” to limit on-target, off-tumor toxicities of chimeric antigen receptor T cells. Front Pharmacol 5, 235–235 (2014).

20. Wu, C.Y., Roybal, K.T., Puchner, E.M., Onuffer, J. & Lim, W.A. Remote control of therapeutic T cells through a small molecule-gated chimeric receptor. Science 350, aab4077 (2015).

21. Rodgers, D.T. et al. Switch-mediated activation and retargeting of CAR-T cells for B-cell malignancies. Proceedings of the National Academy of Sciences 113, E459–E468 (2016).

22. Duong, M.T. et al. Two-Dimensional Regulation of CAR-T Cell Therapy with Orthogonal Switches. Mol Ther Oncolytics 12, 124–137 (2019).

23. Lee, Y.G. et al. Use of a Single CAR T Cell and Several Bispecific Adapters Facilitates Eradication of Multiple Antigenically Different Solid Tumors. Cancer Res 79, 387–396 (2019).

24. Moghimi, B. et al. Preclinical assessment of the efficacy and specificity of GD2-B7H3 SynNotch CAR-T in metastatic neuroblastoma. Nature communications 12, 511 (2021).

25. Rodgers, D.T. et al. Switch-mediated activation and retargeting of CAR-T cells for B-cell malignancies. Proceedings of the National Academy of Sciences of the United States of America 113, E459–468 (2016).

26. Kim, M.S. et al. Redirection of genetically engineered CAR-T cells using bifunctional small molecules. Journal of the American Chemical Society 137, 2832–2835 (2015).

27. Giordano-Attianese, G. et al. A computationally designed chimeric antigen receptor provides a small-molecule safety switch for T-cell therapy. Nat Biotechnol (2020).

28. Raney, K.D., Sharma, S.D., Moustafa, I.M. & Cameron, C.E. Hepatitis C virus non-structural protein 3 (HCV NS3): a multifunctional antiviral target. J Biol Chem 285, 22725–22731 (2010).

29. Mosure, K.W. et al. Preclinical Pharmacokinetics and In Vitro Metabolism of Asunaprevir (BMS-650032), a Potent Hepatitis C Virus NS3 Protease Inhibitor. J Pharm Sci 104, 2813–2823 (2015).

30. Butko, M.T. et al. Fluorescent and photo-oxidizing TimeSTAMP tags track protein fates in light and electron microscopy. Nature neuroscience 15, 1742–1751 (2012).

31. Jacobs, C.L., Badiee, R.K. & Lin, M.Z. StaPLs: versatile genetically encoded modules for engineering drug-inducible proteins. Nat Methods 15, 523–526 (2018).

32. Juillerat, A. et al. Modulation of chimeric antigen receptor surface expression by a small molecule switch. BMC Biotechnol 19, 44 (2019).

33. Yu, S., Yi, M., Qin, S. & Wu, K. Next generation chimeric antigen receptor T cells: safety strategies to overcome toxicity. Mol Cancer 18, 125 (2019).

34. Hong, M., Clubb, J.D. & Chen, Y.Y. Engineering CAR-T Cells for Next-Generation Cancer Therapy. Cancer cell (2020).

## Reference

1. Kennedy, A. & Cribbs, A.P. Production and Concentration of Lentivirus for Transduction of Primary Human T Cells. Methods Mol Biol 1448, 85–93 (2016).

2. Wang, N. et al. Efficacy and safety of CAR19/22 T-cell cocktail therapy in patients with refractory/relapsed B-cell malignancies. Blood 135, 17–27 (2020).

3. Biddison, W.E., Lichtenfels, R., Adibzadeh, M. & Martin, R. Measurement of polyclonal and antigen-specific cytotoxic T cell function. Curr Protoc Immunol (2001).

