## supplemental file for "A Recurring Chemogenetic Switch for Chimeric Antigen Receptor T Cells"

**Supplementary Table 1:** Sequence Information

| **Fragments** | **Sequences** |
| --- | --- |
| Anti-human CD19 scFv | MALPVTALLLPLALLLHAARPIPDIQMTQTTSSLSASLGDRVTISCRASQDISKYLNWYQQKPDGTVKLLIYHTSRLHSGVPSRFSGSGSGTDYSLTISNLEQEDIATYFCQQGNTLPYTFGGGTKLEITGSTSGSGKPGSGEGSTKGEVKLQESGPGLVAPSQSLSVTCTVSGVSLPDYGVSWIRQPPRKGLEWLGVIWGSETTYYNSALKSRLTIIKDNSKSQVFLKMNSLQTDDTAIYYCAKHYYYGGSYAMDYWGQGTSVTV |
| Hinge and cytoplasmic regions | TTTPAPRPPTPAPTIASQPLSLRPEACRPAAGGAVHTRGLDFAFWVLVVVGGVLACYSLLVTVAFIIFWVRSKRSRLLHSDYMNMTPRRPGPTRKHYQPYAPPRDFAAYRSKRGRKKLLYIFKQPFMRPVQTTQEEDGCSCRFPEEEEGGCELRVKFSRSADAPAYQQGQNQLYNELNLGRREEYDVLDKRRGRDPEMGGKPRRKNPQEGLYNELQKDKMAEAYSEIGMKGERRRGKGHDGLYQGLSTATKDTYDALHMQALPPR |
| HCV-NS3 T54A | DEMEECSQHGGSGGSTGCVVIVGRIVLSGSGTSAPITAYAQQTRGLLGCIITSLTGRDKNQVEGEVQIVSTATQTFLATCINGVCWAVYHGAGTRTIASPKGPVIQMYTNVDQDLVGWPAPQGSRSLTPCTCGSSDLYLVTRHADVIPVRRRGDSRGSLLSPRPISYLKGSSGGPLLCPAGHAVGLFRAAVCTRGVAKAVDFIPVENLETTMRSPVFTDNSSPPAVTLTH |
| Firefly luciferase | MEDAKNIKKGPAPFYPLEDGTAGEQLHKAMKRYALVPGTIAFTDAHIEVDITYAEYFEMSVRLAEAMKRYGLNTNHRIVVCSENSLQFFMPVLGALFIGVAVAPANDIYNERELLNSMGISQPTVVFVSKKGLQKILNVQKKLPIIQKIIIMDSKTDYQGFQSMYTFVTSHLPPGFNEYDFVPESFDRDKTIALIMNSSGSTGLPKGVALPHRTACVRFSHARDPIFGNQIIPDTAILSVVPFHHGFGMFTTLGYLICGFRVVLMYRFEEELFLRSLQDYKIQSALLVPTLFSFFAKSTLIDKYDLSNLHEIASGGAPLSKEVGEAVAKRFHLPGIRQGYGLTETTSAILITPEGDDKPGAVGKVVPFFEAKVVDLDTGKTLGVNQRGELCVRGPMIMSGYVNNPEATNALIDKDGWLHSGDIAYWDEDEHFFIVDRLKSLIKYKGYQVAPAELESILLQHPNIFDAGVAGLPDDDAGELPAAVVVLEHGKTMTEKEIVDYVASQVTTAKKLRGGVVFVDEVPKGLTGKLDARKIREILIKAKKGGKIAV |

**Supplementary Table 2:** Primer sequences

| **Primers** | **Sequences** |
| --- | --- |
| CAR19-F1 | GGATCTATTTCCGGTGAATTCGCCACCATGGCGCTGCCTG |
| CAR19-R1 | CCGCGGCGCAGGTGTCGTGGTCACCGTAACGGAGGTTCCTTGTCCCCAAT |
| CAR19-F2 | ATTGGGGACAAGGAACCTCCGTTACGGTGACCACGACACCTGCGCCGCGG |
| CAR19-R2 | AGGTTGATTGTTCCAGACGCGTTTATCTCGGAGGCAGAGCCTGCATATG |
| NS3-F | GAAGCGACGAAATGGAGGAATGTTC |
| NS3-R | AGATCCACCATGGGTGAGAGTGACA |
| Luc-F | CGGAATTCGCCACCATGGAAGACG |
| Luc-R | GCTCTAGATTACACGGCGATCTTTCC |
| Lenti-F | CTGTGACCGCATTGCTCCTT |
| Lenti-R | AGTCCCGTCTGGCTTCTGCT |

**Supplementary Table 3:** Antibodies and providers

| **Antibody** | **Vendor** | **Part number** |
| --- | --- | --- |
| Alexa Fluor 647 rabbit anti-mouse F(ab)^2^ | Jackson ImmunoResearch Laboratories | 315-606-003 |
| FITC human CD19 | AcroBiosystems | CD9-HF251 |
| PE anti-human CD19 | BD Biosciences | 561741 |
| FITC anti-human CD3 | BD Biosciences | 555332 |
| PE anti-human CD4 | BD Biosciences | 555347 |
| APC anti-human CD3 | BioLegend | 300412 |
| APC anti-human CD69 | BioLegend | 310910 |
| PE anti-human CD25 | BioLegend | 302606 |
| FITC anti-human CD4 | BioLegend | 317408 |
| APC anti-human CD8a | BioLegend | 301014 |
| FITC anti-human CD45 | BioLegend | 304006 |
| Annexin V Apoptosis Detection Kit | BD Biosciences | V13241 |
| CFSE Cell Proliferation Kit | BD Biosciences | C34570 |
| PE anti-human CD107a | BD Biosciences | 560948 |
| Protein Transport Inhibitor | BD Biosciences | 554724 |

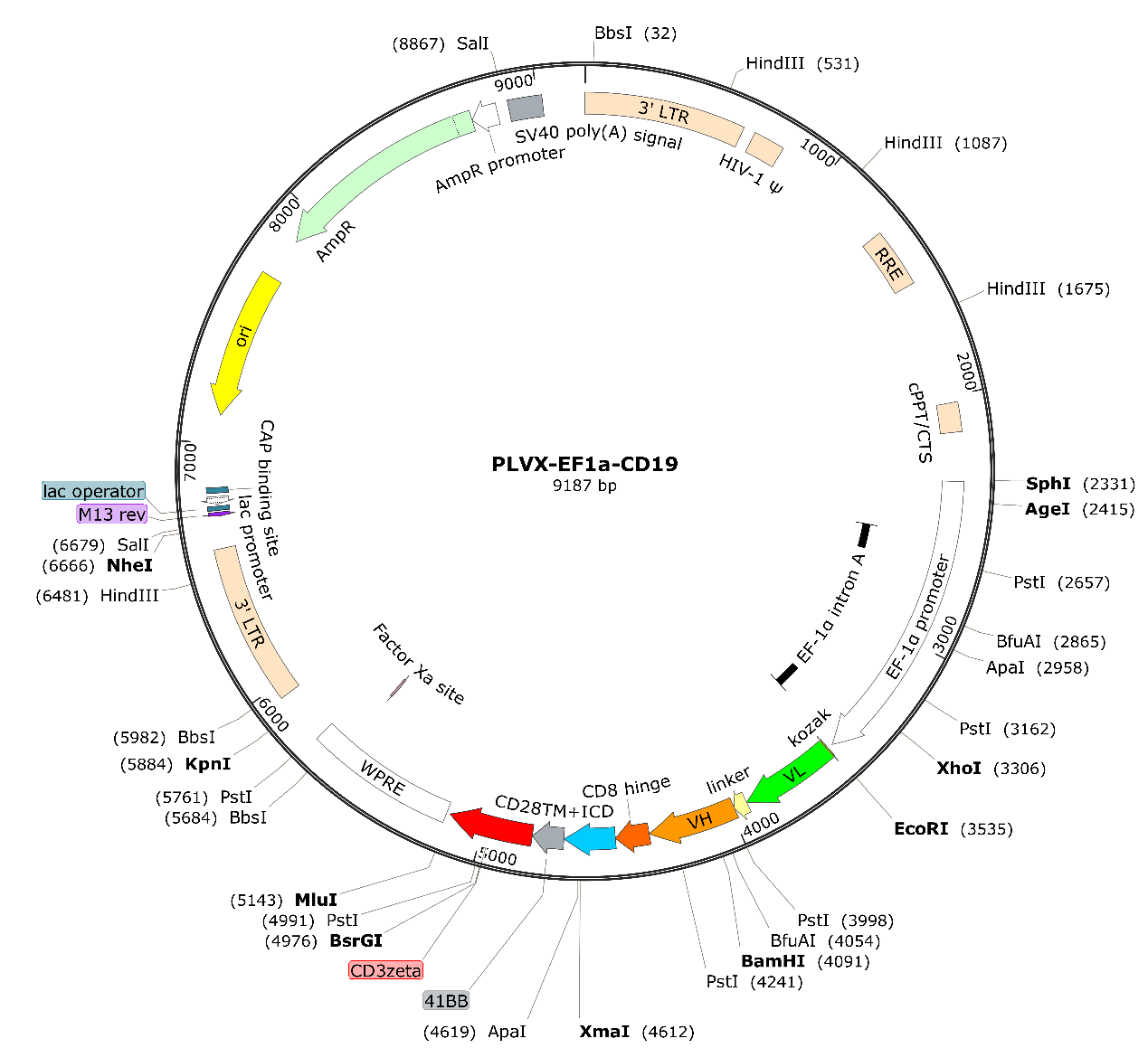

**Supplementary Figure 1.** The plasmid map for pLVX-EF1a-CAR19

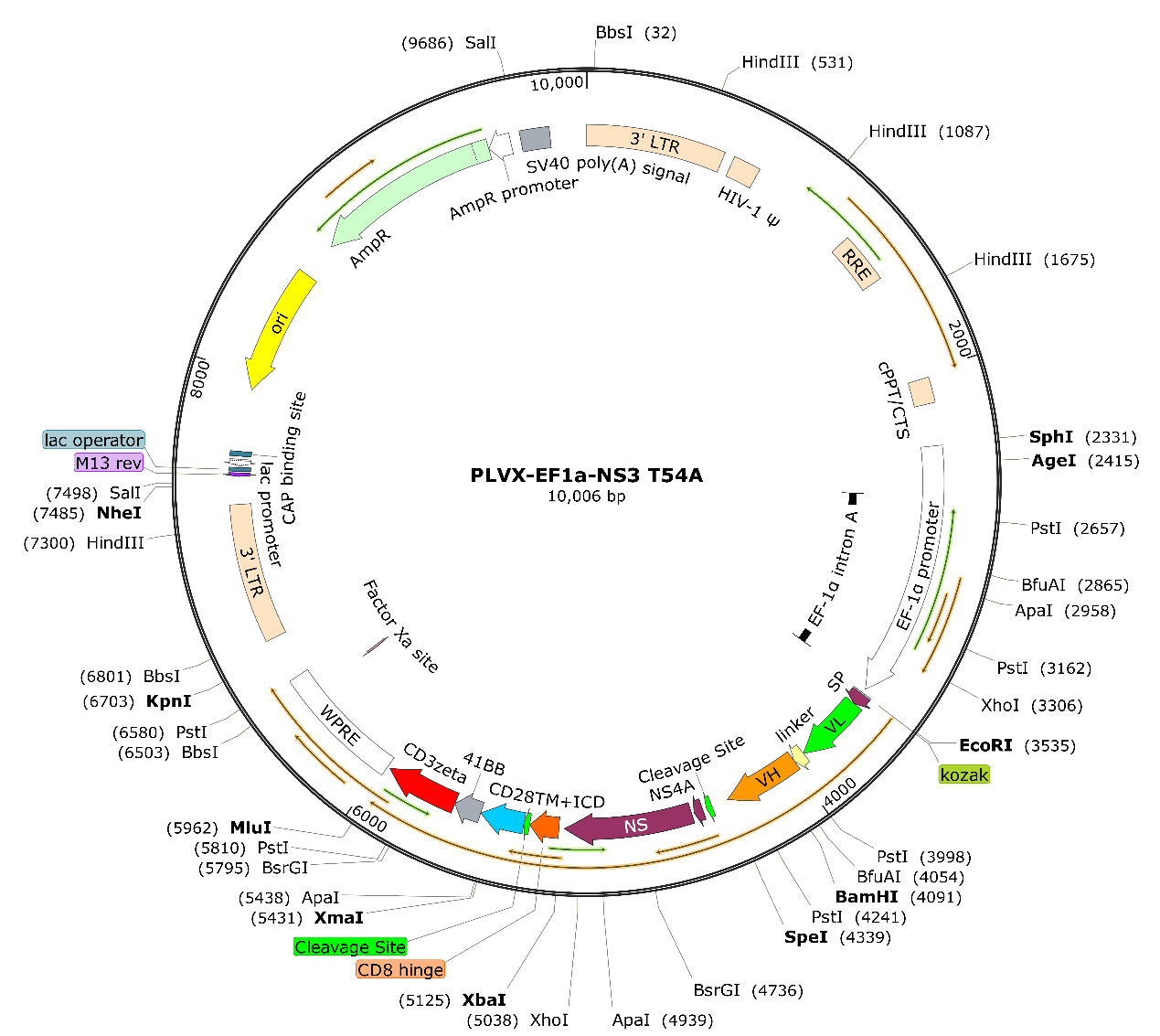

**Supplementary Figure 2.** The plasmid map for pLVX-EF1a-sCAR19

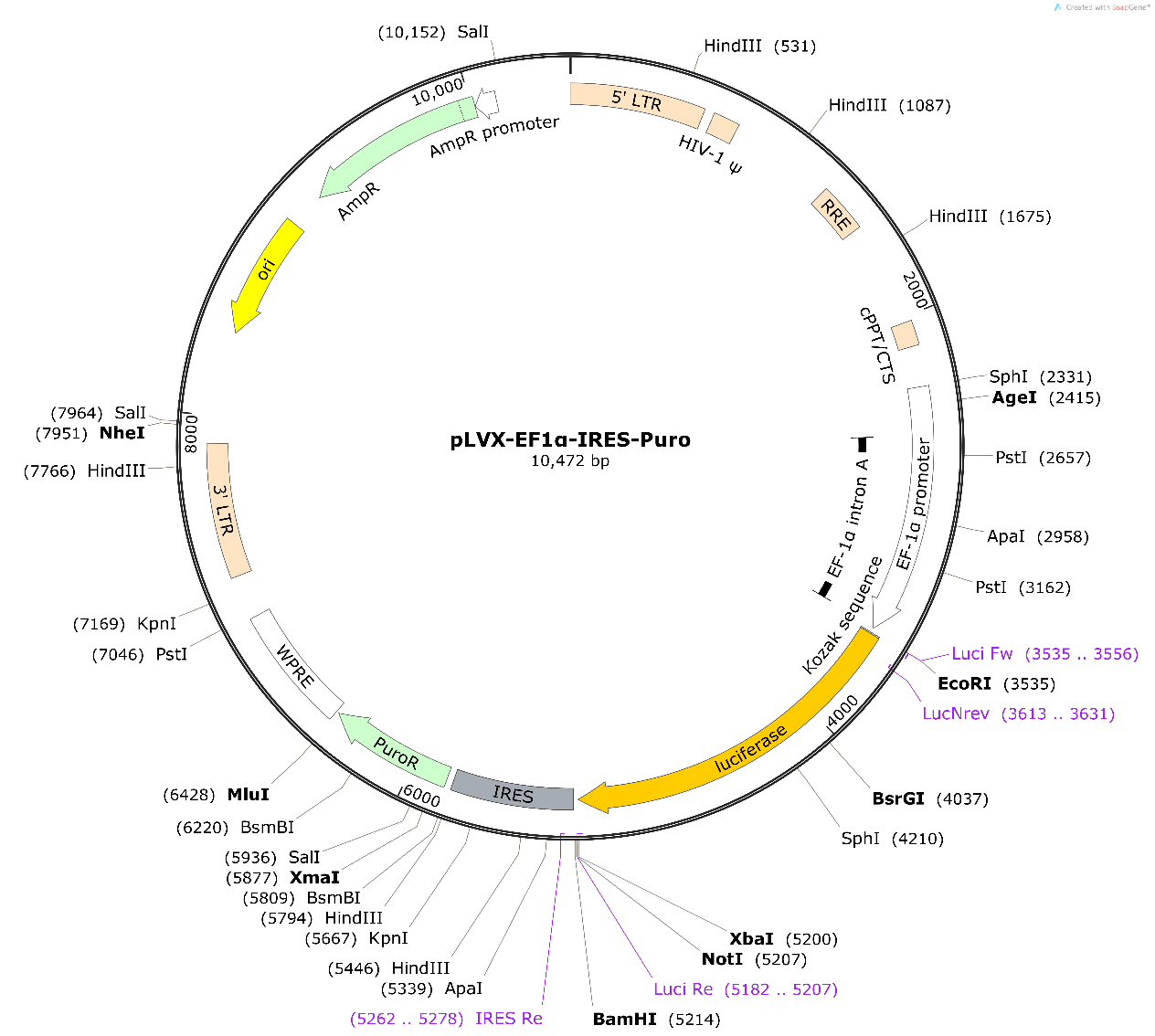

**Supplementary Figure 3.** The plasmid map of pLVX-Luc-Puro

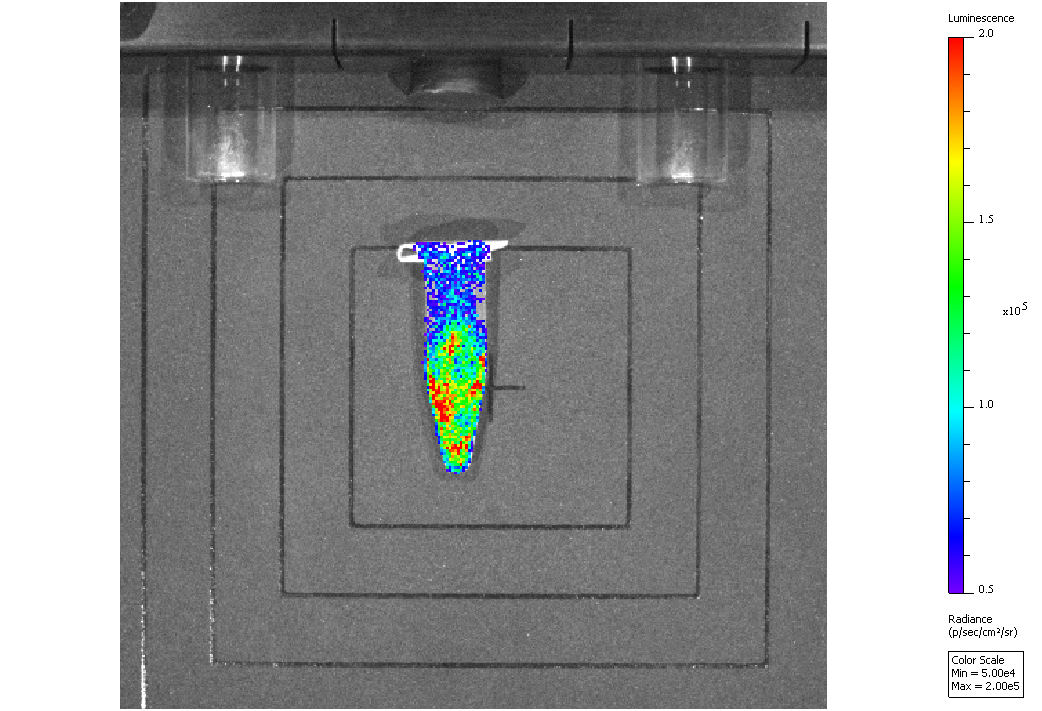

**Supplementary Figure 4.** Raji-Luc Cell imaging
